# Microbial autotrophy is widespread across soils and most prevalent in deep and saturated environments

**DOI:** 10.64898/2026.05.21.726994

**Authors:** Amelia Nelson Kuhn, Alexander L. Jaffe, Petar I. Penev, Kaitlin E. Creamer, Bethany C. Kolody, Preston M. Tasoff, Marcos Voutsinos, Jennifer Pett-Ridge, Jillian F. Banfield

## Abstract

Genes for CO_2_ fixation occur in soil microorganisms, but little is known about the pathways that are most common across ecosystem types, the organisms with these genes, where different CO_2_ fixation pathways are most prevalent, and the energy sources that support autotrophy across ecosystems. Here, we investigated microbial capacity for autotrophy in soils using 853 metagenomes and 201 metatranscriptomes from a wide range of terrestrial ecosystems (agricultural soils, wetlands, weathering rock). Autotrophy-associated RuBisCO (Form I and II) is widely encoded across all soils and occurs in bacteria from numerous lineages (38 phyla). RuBisCO Form IE is consistently more phylogenetically diverse in soils than in marine ecosystems, suggesting that it may have evolved to function in soil-like environments. A newly discovered deeply branching Form I RuBisCO, Form I’’’, supports the hypothesis that Form I RuBisCO originated in anaerobic environments. Further, saturated soils harbor more, and more distinct, autotrophic microbes, many of which may use the Calvin-Benson-Bassham cycle or Wood-Ljungdahl pathway for CO_2_ fixation. Overall, the results indicate that autotrophy is a particularly important metabolism in deep, saturated soils and weathering rock.

## Background

Soils are one of the largest global carbon (C) reservoirs, holding nearly 80% of the C stored in terrestrial ecosystems (∼3100 Gt C(1)). Given the substantial losses of soil C caused by human activity(2) and the increasing concern over climate change, soil C sequestration and stabilization has attracted attention as a promising strategy to mitigate anthropogenic greenhouse gas emissions (3–5). The soil microbiome plays a critical role in the soil C cycle, as microbial growth and activity drive the formation(6–10) and stabilization(11,12) of soil organic matter (SOM). Through these processes, microbes create products that can either remain stored in soils for extended periods or be remineralized, releasing carbon dioxide (CO_2_) back into the atmosphere, thus fundamentally influencing the soil and global C budget(13).

Conceptual frameworks that overview how soil C is formed emphasize the role of microbial decomposers and their interactions with plant-derived C inputs, such as root exudates and leaf litter. Most frameworks highlight soil heterotroph’s interactions with plant-derived C inputs as central to SOM formation (12,14), although the proportion of plant-derived SOM can vary greatly across ecosystems(15,16). These frameworks overlook a possible contributor to soil C: the direct microbial fixation of CO_2_ into microbial biomass, or autotrophy. This process might play a significant role in SOM formation under some conditions. In fact, prior studies using soil incubation experiments have demonstrated active CO_2_ incorporation into microbial biomass via autotrophy (17–19) and suggest that this process could contribute an estimated 0.6-4.9 Pg C per year to soils (20). However, this process has not been investigated in soils broadly, or across ecosystems.

The Calvin-Benson-Bassham (CBB) cycle is the ecologically dominant pathway for C fixation via microbial autotrophy. Its key enzyme, ribulose-1,5 bisphosphate carboxylase/oxygenase (RuBisCO), catalyzes CO_2_ fixation by adding CO_2_ to ribulose-1,5-bisphosphate, forming two molecules of 3-phosphoglycerate (21). RuBisCO Forms I and II are both CBB-associated, but differ in substrate specificity for CO_2_ versus O_2_ and may occupy distinct oxygen niches (21,22). Over the past few years a number of new forms of RuBisCO related to Form Is but basal to Forms IA-IE have been discovered via metagenomic sequence analysis (23). Notably, they lack the small subunit yet have been shown to perform CO_2_ fixation, likely with generally low ability to discriminate between CO_2_ and O_2;_ thus, these new forms have been preferentially identified in anoxic environments (21,23). To date, groups basal to forms IA-E include distinct branches and include Iα, Iβ (previously I’), Iγ (previously I”) and Iδ (24). These findings raise the possibility of additional new forms of RuBisCO that can fix CO_2_, especially in low oxygen or anoxic soils.

Microbially-encoded RuBisCO has been documented in soils across diverse ecosystems, including agricultural fields(19,25–30), grasslands(31–33), wetlands(30,32– 34), forests(17,30,32,35), and deserts(36–39). However, despite these insights, our broader ecological and functional understanding of microbially-encoded RuBisCO in soils remains incomplete. Key knowledge gaps include the RuBisCO distribution across different ecosystems, the taxonomic- and form-level diversity of RuBisCO-encoding microbes in soils, the metabolic contexts in which RuBisCO operates across various terrestrial ecosystems, the role of different RuBisCO-encoding taxa or forms in the generation of persistent soil carbon, and the prominence or coexistence of the CBB cycle in soils with other autotrophic pathways. Addressing these unknowns is critical for understanding the role of microbial autotrophy in soil C cycling and potential shifts in autotrophy in response to climate change.

Here, we leveraged an extensive dataset of 853 soil metagenomes and 201 metatranscriptomes from agricultural soils (e.g., rice paddies, mixed-use dairy farm, and agricultural field), wetlands, grasslands, subalpine forests and river floodplains, and a weathering granite rock profile. Across these ecosystems, we identified 7,140 unique RuBisCO sequences, along with other autotrophic pathway genes, with abundance impacted by soil depth and saturation. CBB-associated RuBisCO forms were most abundant in weathering rock and deep unsaturated mineral soils, while alternate autotrophic pathways (Wood-Ljungdahl and 3-Hydroxypropionate/4-hydroxybutyrate cycle) were dominant in deep, saturated soils. These results indicate that microbial autotrophy is widespread in soils across terrestrial ecosystems and is influenced by soil conditions, with trace gas (H_2_ oxidation) potentially supporting autotrophy in deep soils.

## Methods

### Sample set

To inventory the diversity and distribution of microbial chemoautotrophy in soils, we collated 853 metagenomes and 210 corresponding metatranscriptomes from a suite of soil samples representing various terrestrial ecosystems (e.g., vernal pool, rice paddy, Mediterranean grassland, etc.) and soil depths (≤25 cm classified as shallow, 25-50 cm as mid, >50 cm as deep), hereafter referred to as soil conditions. For analyses presented here, we separated soils into those that are more permanently saturated or were submerged in water when sampled (wetlands, rice paddies) and those that represented ecosystems that are not permanently saturated or often submerged in water (subalpine floodplain and hillslope, agricultural fields, grasslands). Sample metadata, including soil taxonomy classifications from the USDA Web Soil Survey, and corresponding publications are detailed in **Supplementary Data 1** (Sheet A).

### Annotation and curation of soil-derived RuBisCO sequences

Reads were all assembled into annotated scaffolds following the ggKbase metagenomic data preparation pipeline (see https://ggkbase-help.berkeley.edu/overview/data-preparation-metagenome/ for details). Proteins were predicted from assembled metagenomic scaffolds using Prodigal (V2.6.3) and filtered to those on contigs >1000bp long. All protein sequences were combined and searched against a series of hidden Markov models (HMMs)(40) made using sequences reported in Prywes et al.(21) describing the major forms within the RuBisCO superfamily using HMMER (v3.3) (41). Hits were filtered to those with a score >100 and covered at least 50% of the model. If a query sequence hit multiple HMMs following this filtering, the model with the highest score was chosen. These RuBisco form-level annotations were confirmed phylogenetically using graftM (v0.13.1)(42). To aid further curation, sequences were clustered at 95% identity using vsearch (v2.13.3)(43) and, in cases where cluster sequences HMM and graftM annotations differed, we used phylogenetic tree visualization for manual curation and verification (189 clusters encompassing 430 protein sequences). Briefly, sequences were aligned using MAFFT(44)(v7.505) and the resultant tree was built using FastTree(45)(v2.1.11) and visualized with iTol(46). Final form-level annotations were chosen depending on their position relative to the reference sequences in the tree. Sequences were dereplicated using mmseqs (--min-seq-id 1.0)(47) and filtered to those >400 amino acids long to remove fragmented proteins, resulting in 7,140 unique RuBisCO protein sequences (pipeline overview in **Fig. S1**; sequence details in **Supplementary Data 1**, Sheet C). RuBisCO Form I subtypes were further annotated using phylogenetic tree visualization with Form I reference sequences from Pyrwes et al.(21) and Ray et al.(37). RuBisCO taxonomy was determined through a voting scheme in which taxonomy was assigned via comparison with a sequence database (protein annotations in UniProt(48) and ggKbase: https://ggkbase.berkeley.edu/) when the same taxonomic assignment received greater than 50% votes.

### Alternate autotrophic pathways and metabolic annotation in metagenomes

In addition to RuBisCO, alternative autotrophic pathways and supporting metabolic genes were annotated from the metagenomes to identify the prevalence of autotrophic metabolisms distinct from CBB and catabolic processes that potentially energetically support the CBB cycle in soils. For alternate autotrophic pathways, genes associated with the Wood-Ljungdahl pathway (WL), 3-Hydroxypropionate cycle (3HP), 3-Hydroxypropionate/4-hydroxybutyrate cycle (3HP/4HB), and the reductive tricarboxylic acid cycle (rTCA) were annotated. To inventory supporting catabolisms, we identified protein sequences associated with nitrogen (N), sulfur (S), hydrogen (H_2_), and iron (Fe) oxidative pathways. Using hmmsearch, we screened all sequences against a set of HMMs with predefined thresholds (49–51). In some instances, results were further refined through manual curation and phylogenetic analyses.

For ammonia monooxygenase (amoA; K10944), we validated sequences annotated via HMMs by aligning them with a diverse set of reference sequences(52) using MAFFT(44)(v7.505). A phylogenetic tree was constructed with FastTree(45)(v2.1.11) and visualized with iTol(46), and hits clustering with pmoA reference sequences in the resultant tree were discarded. Similarly, nitrite oxidoreductase (K00370) sequences were validated using reference sequences from UniProt(48). For putative dissimilatory sulfite reductase (dsr) genes, we compared protein sequences to reference sequences (53) using blastp. Final dsr genes retained had at least 50% identity to members of the oxidative clade. NiFe hydrogenases involved in aerobic H_2_ oxidation were annotated following methods detailed in (54). Briefly, sequences were aligned against a set of NiFe hydrogenase reference proteins (51) using DIAMOND (v2.1.9.63)(55). Hits were filtered based on the following criteria: bit score >100, query or subject coverage ≥80%, minimum identity score ≥60%, and classification into energy-yielding forms 1d, 1l, or 2a. Annotations were reconfirmed through phylogenetic analysis via tree-building with the reference proteins. Iron oxidation genes were annotated using HMMs and trusted cutoff scores provided by FeGenie(50). Other hydrogenases were annotated using HYDdb(51). Final annotated genes were dereplicated using mmseqs (-- min-seq-id 1.0)(47). All gene names, associated accession numbers, annotation techniques and, if relevant, cutoff thresholds used are detailed in the **Supplementary Data 1** (Sheet B). The final dataset comprised 483,076 protein sequences, detailed in **Supplementary Data 1** (Sheet D).

### Metagenomic and metatranscriptomic read mapping to the gene dataset

Trimmed metagenomic reads were mapped to the final gene database nucleic acid sequences using Bowtie2 (v2.5.4)(56) with default parameters. Gene coverage across samples was calculated using coverM contig (v0.7.0)(57) with the “Trimmed Mean” (hereafter, referred to as ‘TMM’) method, and final reported TMM value was normalized by sequencing depth of the associated sample metagenome (Gbp). Trimmed metatranscriptomic reads were mapped to the gene database nucleic acid sequences using Bowtie2 (v2.5.4)(56) and filtered to 99% identity. Mapped reads were further filtered to exclude unmapped reads and retain only those with a mapping quality (MAPQ) ≥20 and counts were generated using featureCounts (v2.0.6)(58). Final transcript abundances were normalized by dividing read counts by gene length. Normalized gene TMM and transcript abundance of genes across samples are included in the **Supplementary Data 2** and **Supplementary Data 3**. To ensure that spurious results were not resulting from mapping metatranscriptomic reads to genes recovered from all samples (instead of mapping to only samples recovered from that project or field site), we tested with read recruitment with a subset of samples from the vernal pool sites to ensure similar trends (see **Supplementary Text**).

### Gene coverage analyses and statistics

To identify how the autotrophic soil microbiome (via metagenomic and metatranscriptomic read mapping) differed across ecosystem type and soil depth, statistical analyses were performed using R (v4.4.2)(59) with significance values accepted at *p*<0.05. Differences in gene coverage and RuBisCO-encoding community diversity were assessed and tested using pairwise Wilcoxon signed-rank tests with a Bonferroni p-value adjustment using the function “stat_compare_means” in the ggpubr R package(60). Spearman tests were used to test the strength of correlation between the gene coverage of a given function and soil pH across different soil depths using the “cor.test” function in the stats core R package(59). To visualize beta diversity of soil microbial communities encoding genes associated with different autotrophy pathways, non-metric multidimensional scaling (NMDS) was conducted using Bray-Curtis dissimilarity in the vegan(61) and phyloseq(62) R packages. To statistically test what was driving the beta diversity, nonparametric permutational multivariate analysis of difference (PERMANOVA)(63) was performed against the Bray-Curtis dissimilarity matrices using the vegan(61) “adonis2” function. The R package ggplot2(64) or iTOL(46) were used to make all plots, which were edited and made more clear for readers (e.g., making outlines larger on points to increase visibility) in Adobe Illustrator (v29.5).

## Results

### RuBisCO is widely detected in weathering rock and diverse soil microbiomes

A total of 853 soil metagenomes and 201 soil metatranscriptomes were used to inventory the distribution and diversity of RuBisCO-encoding and autotrophic soil microbes. Ecosystems in the sample set included grasslands(65–68), rice paddies (Kolody et al., in prep), subalpine pine forests (70, 71, 72, Tasoff et al., in prep), mixed-use agricultural soils (Creamer et al., in prep), wetlands(72–74), and weathering granite rock(75), providing a range of soil abiotic conditions and saturated and unsaturated soils (**Supplementary Data 1**, Sheet A; **Table S1**). We identified 7,140 dereplicated, phylogenetically verified RuBisCO and RuBisCO-like sequences from 706 of the metagenomes (**Fig. 1**). Although not all metagenomes had RuBisCO in the final curated, unique dataset, all ecosystem types were represented (**Fig. 1**).

**Fig. 1.**
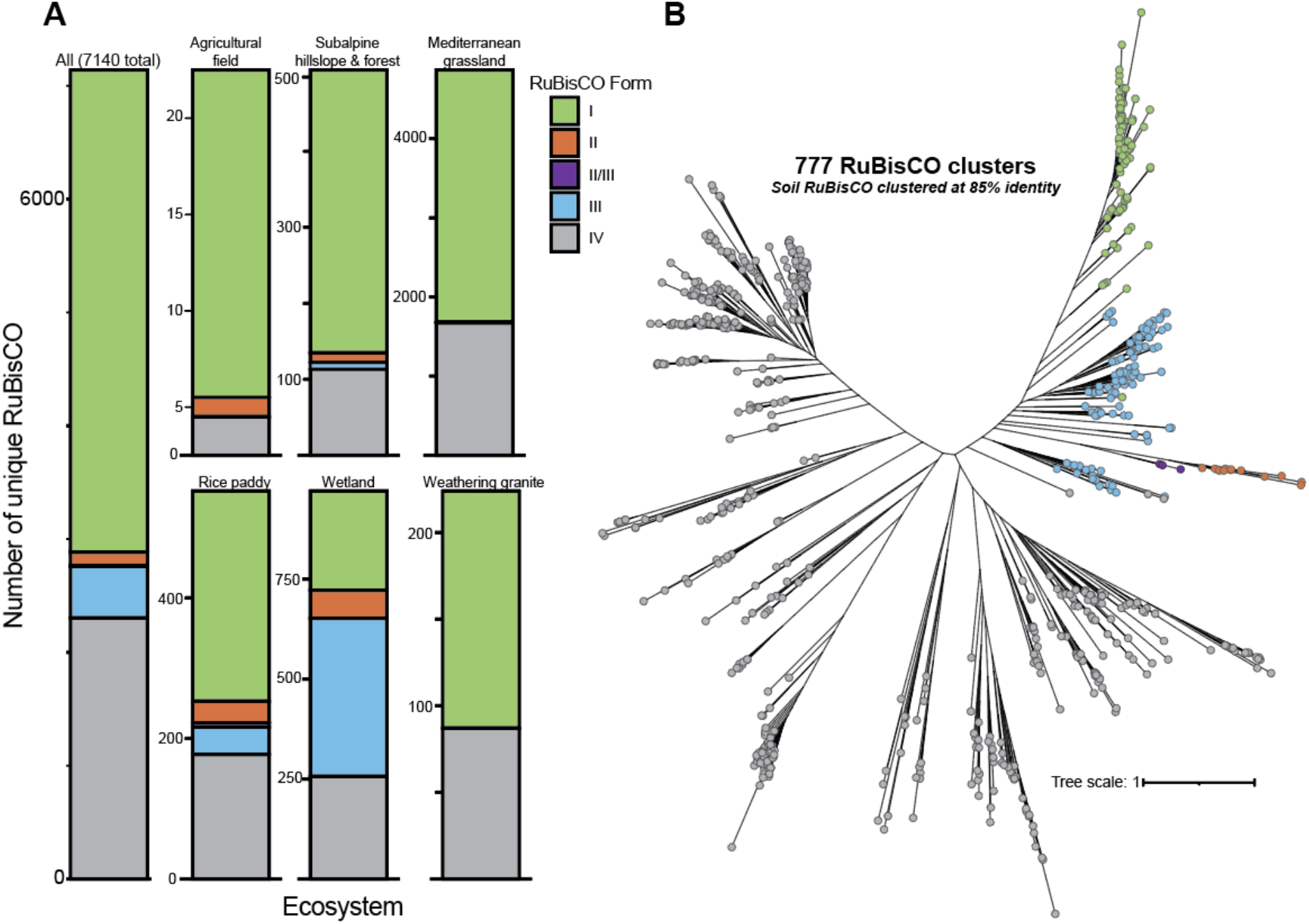
RuBisCO form and phylogenetic diversity across soils. **(A)** Number of unique RuBisCO clusters found in soils from across ecosystems, annotated by form. Ecosystems include an agricultural field, subalpine hillslope and forest (East River, CO, USA; pine and aspen), a mediterranean grassland (CA, USA), rice paddy, wetland sediment (CA, USA), and a weathering rock profile (Ararat, VIC, AUS). **(B)** Concatenated RuBisCO protein phylogeny of all unique RuBisCO found in this study clustered at 85% identity.

### RuBisCO form and abundance vary with soil depth and saturation

Of the unique RuBisCO sequences, 4257 were classified as Form I, 116 as Form II, 6 as Form II/III, 456 as Form III, and 2305 as Form IV. Form IV are RuBisCO-like proteins that are unlikely to function in CO_2_ fixation, so they were not considered further. Soils from constantly saturated ecosystems (e.g., wetland, rice paddies) encoded most of the detected Form II (87%) and Form III (95%) RuBisCO (**Fig. S2**), while unsaturated soils harbored the majority of the Form I RuBisCO (3709 of 4257; **Fig. 1**). Of the Form I sequences, 3870 could be confidently classified into types A through E or other deep branching forms. The majority of the soil-derived Form I RuBisCO were Form IE (1820 sequences) or Form IC (1669). Of the remaining sequences classified as RuBisCO, 180 were Form ID, 83 were Form IB, and 73 were Form IA. An additional 36 sequences were classified as Form 1α, and 9 as Form I’ (**Fig. S3**); these recently described deep branching groups lack the small RuBisCo subunit. The majority of the Form 1α sequences (31 of 36) came from saturated soils (**Fig. S3**).

RuBisCO forms varied in abundance across soil types and depth; the relative abundance of CBB-associated Form I and II genes increased with depth in all soils (**Fig. 2**). However, the highest abundances (normalized by sample sequencing depth) were detected in samples collected from weathering granite (saprolite), material generally found at the base of soil horizons and represent the soil’s parent material (**Fig. 2, Fig. S4**). Mirroring the gene coverage profiles of CBB-associated RuBisCO, phosphoribulokinase (*prk*), another key enzyme in the CBB cycle, increased significantly (*p*<0.0001) with soil depth in nonsaturated soils; the highest gene coverage was also found in samples collected from weathering granite (**Fig. S5**). RuBisCO forms II and III were largely derived from saturated soil metagenomes (**Fig. S2**), and were most abundant in deep (>50 cm) saturated soils (**Fig. S4**).

**Fig. 2.**
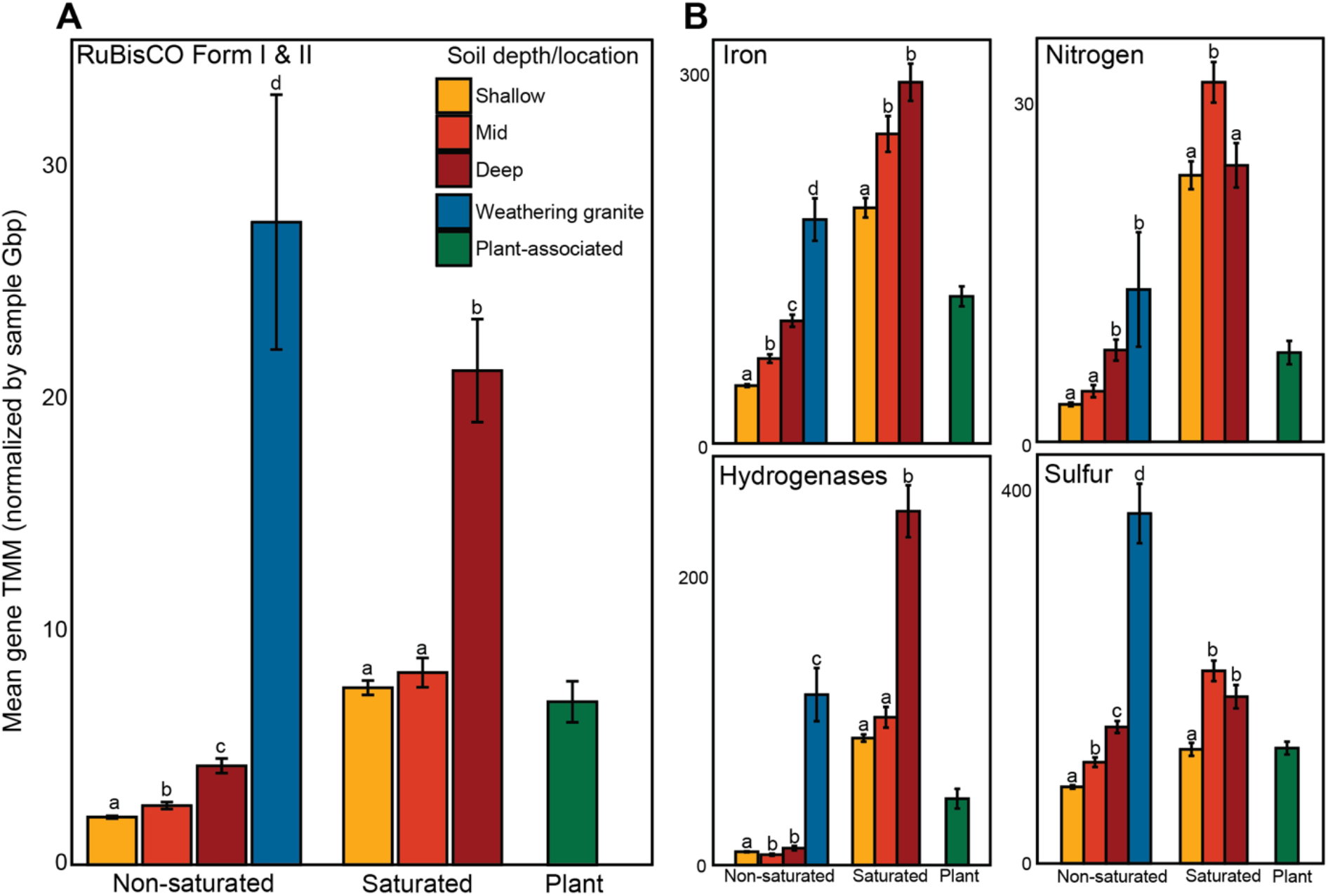
Abundance of soil autotrophy and supportive catabolism genes shift with ecosystem and soil depth. **(A)** Mean sample gene normalized abundance (TMM) of RuBisCO form I & II (summed) across ecosystem type and soil depth (shallow = ≤25 cm, mid = 25-50 cm, deep = >50 cm, granite indicates a weathering granite profile, plant-associated samples were separated by root depth). **(B)** Mean sample gene normalized abundance (TMM) of genes for iron, nitrogen, and sulfur oxidation, and hydrogenases. Bars represent the standard error of the mean. Shared letters represent groups that are not significantly different from one another within ecosystem types, using a Wilcoxon rank-sum test.

### Novel deeply branching Form I RuBisCOs

Included in our dataset were 7 closely related sequences that form a phylogenetic clade distinct from other Form I groups and is basal to the forms IA-E, as well as to I’ and I’’ (**Fig. 3**; navy blue clade). We assigned these 8 sequences to a new clade, which we refer to as Form I’’’. The I’’’ clade is quite distinct from the recently reported Iδ clade(24), which appears to be the deepest branch in the Form I radiation. The I’’’ large subunit RuBisCOs are encoded adjacent to *prk* but no co-encoded small subunit, paralleling findings for all groups basal to forms IA-IE RuBisCOs, which do not require the RuBisCO small subunit to be functional. These novel RuBisCO sequences are derived from rice paddy soils and were all encoded on Chloroflexi genome fragments.

**Fig. 3.**
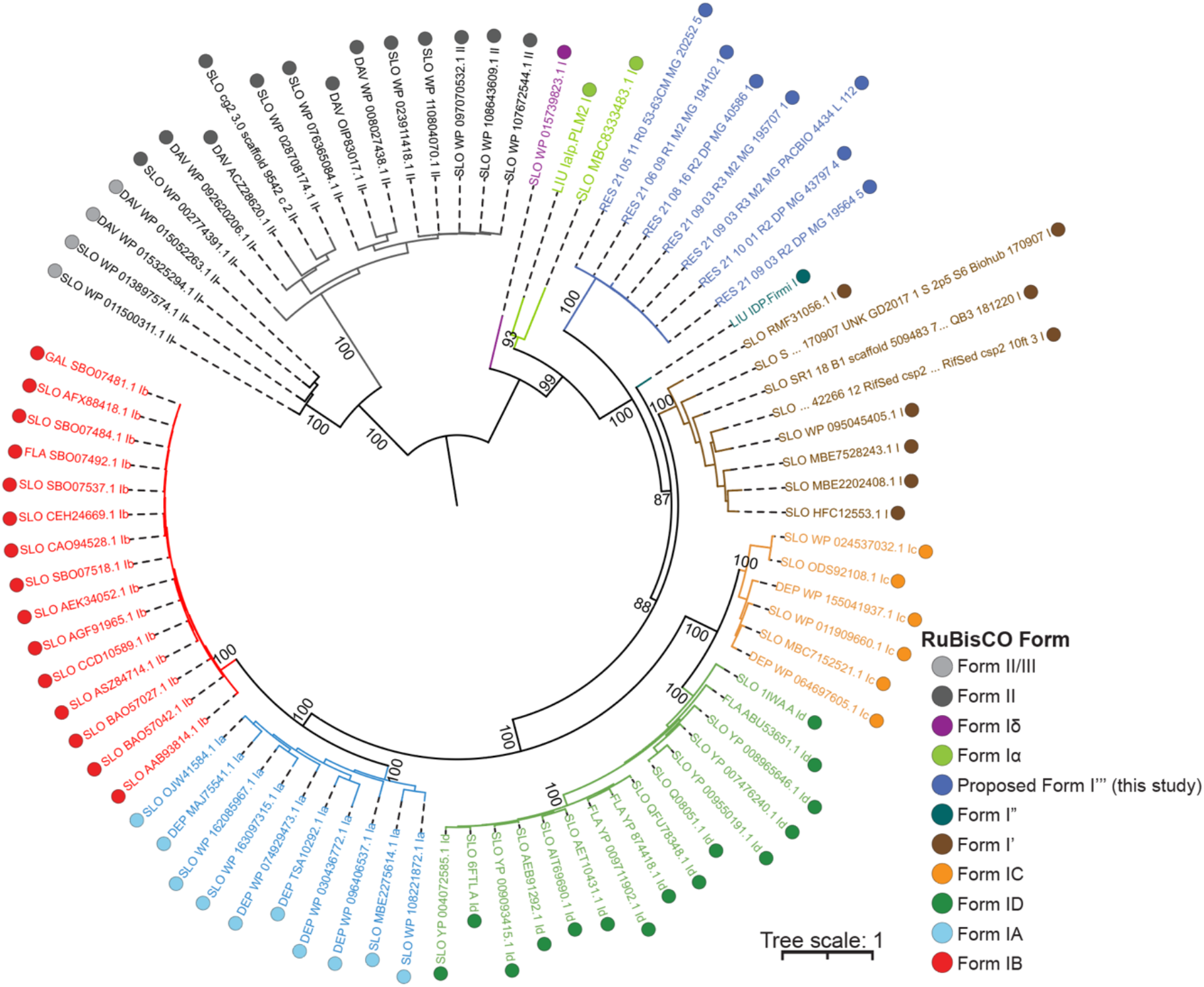
New phylogenetically-diverging Form I RuBisCO. Concatenated RuBisCO protein phylogeny of new deeply branching Form I RuBisCOs with reference sequences from Kehl et al. (2025) (76). Branches annotated as sequence RuBisCO form. Bootstrap values are displayed for nodes defining major RuBisCO Form I subforms; support values for terminal within-subform branching are omitted for clarity.

### Diverse bacteria encode RuBisCO

The Proteobacteria contributed the majority of RuBisCO sequences in our dataset (**Fig. S6**; **Table S2**), with the Alphaproteobacteria (707) and Betaproteobacteria (558) encoding many of the RuBisCO Forms I and II (**Supplementary Data 1**, Sheet C). The majority of CBB-associated RuBisCO were associated with Proteobacteria in saturated soils and Actinobacteria in non-saturated soils (**Fig. S7**). Proteobacteria were the most diverse CBB-associated RuBisCO-encoding group, with 214 distinct lineages encoding Forms I and II across soils (**Fig. S8**). In non-saturated soils, the diversity of organisms with CBB-associated RuBisCO increased with soil depth in non-saturated soils and was significantly higher in non-saturated than saturated soils (**Fig. S9**).

### Non-CBB autotrophy pathways are widely encoded and vary by system type

In addition to RuBisCO, we also inventoried the metagenomic dataset for genes encoding alternate autotrophic pathways, including the Wood-Ljungdahl pathway (WL), 3-hydroxypropionate cycle (3HP), 3-hydroxypropionate/4-hydroxybutyrate cycle (3HP/4HB), and the reductive tricarboxylic acid cycle (rTCA) (**Supplementary data 1**, Sheet B). All inventoried genes were detected in the soil metagenomes (49,675 total dereplicated genes) with the largest number of detected unique genes in the WL and 3HP/4HB pathways.

Saturated soils had higher coverage of genes associated with the WL pathway and 3-HP/4-HB cycle relative to non-saturated soils, with highest coverage in deep saturated soils (**Fig. S5**). Across both saturated and non-saturated soils, genes encoding for the rTCA cycle increased with depth but were completely absent in weathering granite (**Fig. S5**). Genes for the 3-HP cycle were the most enriched in weathering granite samples relative to bulk soils (**Fig. S5**).

In addition to soil conditions (non-saturated vs. saturated) and depth, relative abundances of different autotrophy genes varied strongly by soil type (**Fig. S10**). Ordination analysis of gene coverage profiles for all autotrophy pathways (using Bray-Curtis dissimilarity) revealed that beta diversity was primarily structured by soil condition (PERMANOVA R^2^ = 0.122, *p* = 0.001), with soil depth contributing to a lesser extent (PERMANOVA R^2^ = 0.044, *p* = 0.001; **Fig. S10**). Clustering also broadly aligned with soil saturation: non-saturated ecosystems (weathering granite, grassland, agricultural field, and subalpine soils) clustered along the left side of NMDS1, while continuously saturated ecosystems (rice paddy, wetland) clustered on the right (**Fig. S10**). The subalpine soils that grouped closer to saturated ecosystems originated from the East River floodplain(70), areas that are saturated during high flow following snowmelt. These patterns were not clearly associated with soil pH (**Fig. S11**).

### Catabolic potential for chemoautotrophy differs by soil saturation and depth

Soil microorganisms are often characterized as mainly and unlikely to fix C. Thus, we sought evidence of other metabolic pathways (sulfur, iron, hydrogen, and nitrogen oxidation) that might be linked to autotrophic growth in soils (**Supplementary data 1**, Sheet B). We identified 22,718 unique hydrogenases, 119,551 genes for iron oxidation, 5401 nitrification genes, and 278,759 genes associated with sulfur oxidation. Normalized gene abundances varied significantly by ecosystem and soil condition, with sulfur oxidation genes most abundant in the weathering rock profile, and hydrogenases enriched in both the weathering rock profile and constantly saturated soils (e.g., rice paddy and wetland soils; **Fig. S12**). In non-saturated soils, gene abundances generally increased with depth (**Fig. 2**). In saturated soils, abundance of hydrogenases and iron oxidation genes also increased with depth (**Fig. 2**). Across all pathways, gene abundances for chemoautotrophy were consistently higher in saturated than in non-saturated soils (**Fig. S13**).

### Evidence of active autotrophic communities across soils

To quantify transcript abundance of chemoautotrophic pathways across soils, we mapped available metatranscriptomic reads that accompanied the analyzed metagenomes to the gene dataset (including RuBisCO, alternative autotrophy pathways, catabolisms, hydrogenases). Of the 483,076 genes, 90,900 had 1 or more transcripts mapping to them (∼18.8% of genes) and 2229 of the 7140 RuBisCO had mapped transcripts (∼31% of RuBisCO). Across the RuBisCO forms, 39% of recovered Form I had mapped transcripts (1685 of 4245), 46% of Form II (54 of 116), 0% of Form II/III, 15% of Form III (72 of 456) and 18% of Form IV (418 of 2229). Although the metatranscriptomes were mapped to genes recovered from all samples, most transcripts mapped to genes that were recovered from the same field site (**Table S3, Supplementary text**).

RuBisCO Form I and *prk* were expressed across all sample types, with the highest normalized transcript abundance for CBB in mid-depth saturated soils (25-50 cm depth, **Fig. 4, Fig. S14**). Alternative autotrophy pathway genes (WL, 3HP, 3HP/4HB, and rTCA) also all recruited transcripts, with transcripts for WL most abundant across soils (**Fig. 4, Fig. S15**). Nitrification genes recruited the most transcripts in non-saturated soils, with a clear increase of *amoA* transcript abundance with depth (**Fig. S16**). Transcripts mapping to sulfur oxidation genes (e.g., *soxB, soxY, soxC, fccB*) were more abundant in non-saturated soils and increased with soil depth (**Fig. 4, Fig. S17**). Lastly, hydrogenases likely to permit energy gain via aerobic respiration of H_2_ recruited transcripts across all soil types (**Fig. S19**), although the [NiFe] Hydrogenases (Group 2a) showed no evidence of expression in deep saturated and non-saturated soils (**Fig. 4**).

**Fig. 4.**
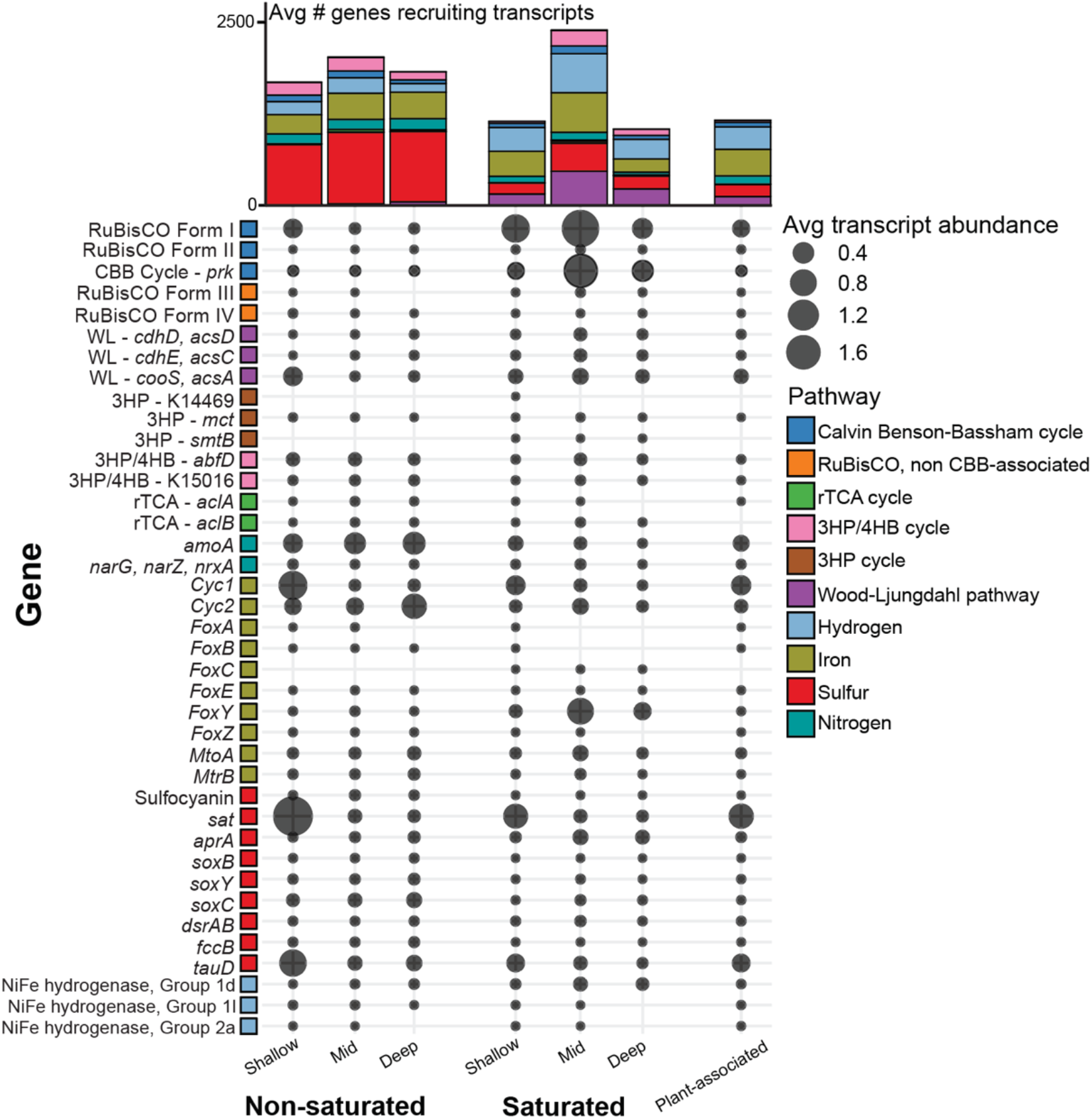
Active autotrophic pathways and metabolic pathways that may support them. Normalized average transcript abundance of genes across different soil types (permanently saturated vs. non-saturated ecosystems) and depths (shallow = ≤25 cm, mid = 25-50 cm, deep = >50 cm) colored by metabolic categories. Hydrogen genes include hydrogenases involved in aerobic H_2_ oxidation and iron, sulfur, and nitrogen genes refer to those involved in their respective oxidative cycles. Saturated samples include those from rice paddies and wetlands and non-saturated includes all ecosystems not permanently saturated (grasslands, agricultural field, subalpine forest, weathering granite). Above, the average number of genes in each metabolic category that recruit transcripts across soil types. Individual sample data points are included in Fig. S14-S18.

### RuBisCO sequence diversity and partitioning between soil and marine environments

To investigate ecosystem-specific selection and variability within the RuBisCO-encoding microbiome, we compared our soil-derived RuBisCO dataset with a published collection of marine RuBisCO sequences from Jaffe et al. (2024)(40). The marine dataset comprised 1,245 non-redundant RuBisCO sourced from 28 open ocean metagenomes spanning depths of 50 to 4,000 meters and public ocean metagenome-assembled genome (MAG) datasets. Following dereplication at 100% amino acid identity, the soil-derived and marine-derived RuBisCO sets showed no sequence overlap, suggesting strong ecosystem-specific partitioning of RuBisCO sequences.

Soil-derived RuBisCO exhibited greater sequence diversity than those from marine environments (**Fig. 5C**). The only exception was Form II/III, which showed higher sequence diversity in the marine dataset; however, this was based on few sequences (14 marine, 6 soil) and may not reflect broader diversity trends. Clustering all 8,377 unique soil and marine RuBisCO sequences together at 80% identity revealed that most clusters were ecosystem-specific: 805 clusters contained only soil-derived sequences, and 256 contained only marine-derived sequences (**Fig. 5**). The largest soil-only cluster (Cluster 123) consisted of 312 RuBisCO sequences, all annotated as Form I and phylogenetically most like Form IE (**Fig. S20**). The largest mixed-origin cluster (Cluster 3), containing 98 marine and 660 soil, was Form IC RuBisCO (**Fig. S20c**).

**Fig. 5.**
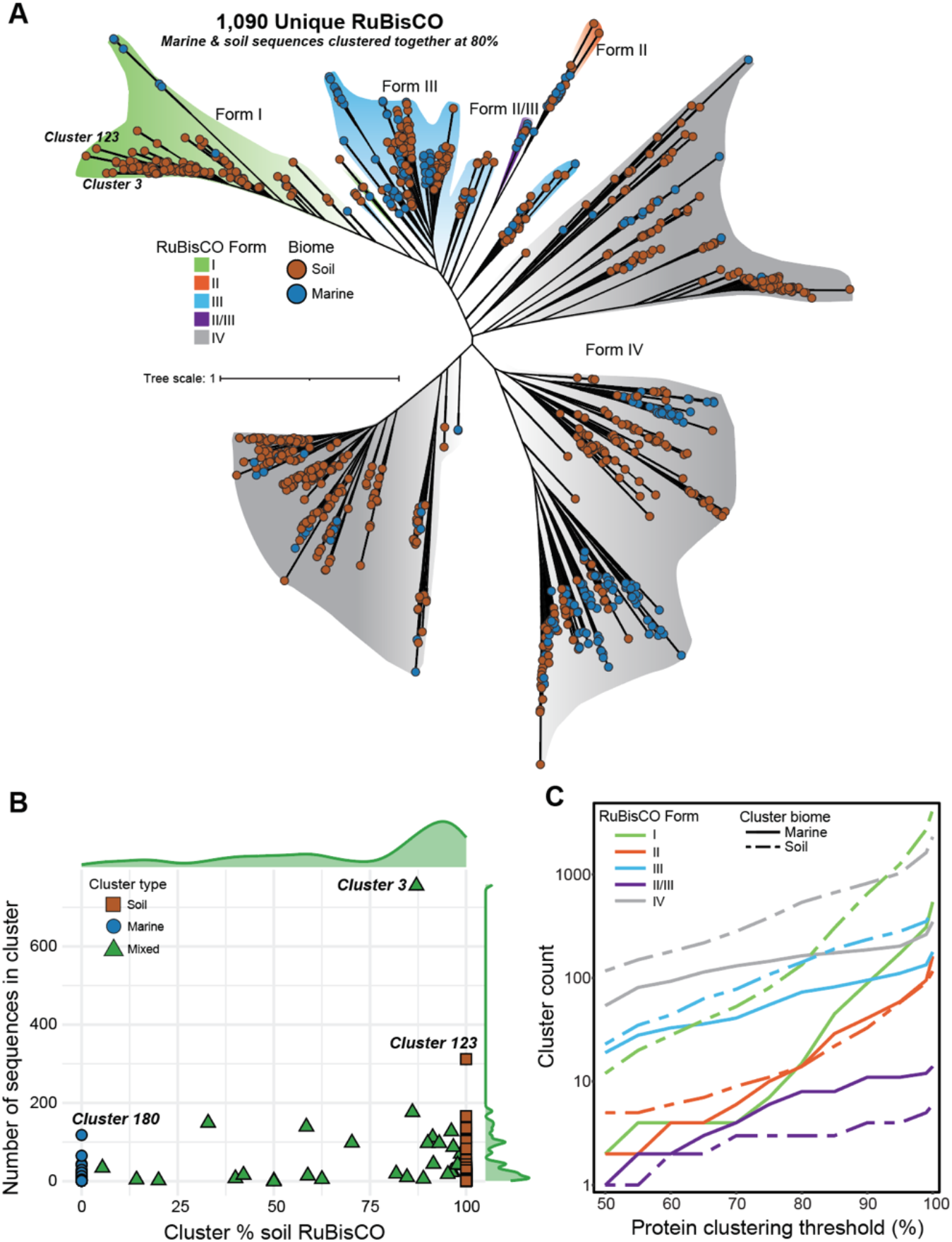
Soil and marine-derived RuBisCO comparative analysis. **(A)** Concatenated RuBisCO protein phylogeny of all unique marine and soil RuBisCO clustered together at 80% identity. Branches annotated as RuBisCO form and labeled with sequence origin biome. **(B)** Information on clusters resulting from clustering all marine and soil sequences together at 80% identity. Points represent each individual resultant cluster, with the x-axis indicating the % of soil RuBisCO sequences making up the cluster and y-axis showing overall size of cluster with corresponding density plots. Points are colored by whether the cluster consists of all soil sequences, all marine sequences, or a mix of the two biomes. **(C)** Line plot showing number of clusters resulting from independently clustering marine and soil RuBisCO sequences at different clustering thresholds (x-axis) and number of total resultant clusters (y-axis). Line colors indicate consensus RuBisCO form of cluster and line stroke indicates biome.

## Discussion

In this study, we provide metagenomic evidence that genes for microbial autotrophy are widely encoded within soil microbiomes. Our meta-analysis of soil metatranscriptomes revealed that RuBisCO is expressed in bacteria from many lineages in soil and weathered rock microbiomes. The increased abundance of CBB-associated RuBisCO and phosphoribulokinase (*prk*) with soil depth and in weathering granite (**Fig. S4**; **Fig. S5**) suggests that autotrophy may be particularly important in environments with limited organic C inputs. The CBB-encoding microbiome was more diverse in non-saturated than saturated soils, regardless of soil depth, likely due to more biodiversity in non-saturated compared to saturated soils (**Fig. S9**). Additionally, autotrophic pathways varied across soil types, with saturated soils selecting for organisms that use the Wood-Ljungdahl pathway (WL) and the 3HP/4HB cycle and weathering rock favoring organisms encoding CBB and 3HP pathways (**Fig. S5**). The abundance of autotrophy genes was not correlated with soil pH (**Fig. S11**) and may instead be driven by other soil properties not measured here (e.g., CO_2_ concentration or organic C availability) or variations in aboveground plant inputs. Prior surveys of deep soil metagenomes, including a comprehensive Critical Zone Observatory-wide study (77), reported no evidence of autotrophy, likely due in part to the absence of a targeted detection and curation pipeline for RuBisCO and other autotrophic pathways. By applying computational pipelines specifically designed to identify RuBisCO and other autotrophic pathway genes, we show here that autotrophic metabolisms are likely widespread across the deep soil microbiome.

The discovery of a new clade of RuBisCOs (**Fig. 3**) that apparently function in the CBB pathway but lack the small subunit increases the diversity of single subunit RuBisCO clades basal to the classic forms IA-IE RuBisCOs (all of which require a small subunit). Prior work on the first discovered clade of RuBisCOs without a small subunit experimentally established their ability to fix CO_2_(23). These sequences (Form I’), along with the new Form I’’’ reported here, are derived from Chloroflexi. As the exact phylogenetic placement, thus branching order in the deep clades, remains uncertain, it is unclear in which bacterial group Form I’s may have originated. The general association of the deep branching single subunit RuBisCOs with anaerobic environments, exemplified by deep wetland soil (e.g., some Iα) and deep rice paddy soil (e.g., all the putative I’’’), support origination of Form I RuBisCO in anaerobic environments where oxygen sensitivity is less detrimental to enzyme function.

RuBisCO-mediated autotrophy is widespread across the deep ocean and largely taxonomically constrained to several orders in the Gammaproteobacteria and SAR324(40). Here, we found more taxonomic and sequence diversity in RuBisCO in soils as compared to marine ecosystems (**Fig. 5C, Fig. S7**), likely reflecting the higher overall microbial and functional diversity typically observed in soils compared to marine ecosystems(78). When soil and marine-derived RuBisCO sequences were combined and clustered together, the largest soil-only RuBisCO cluster was composed of Form IE sequences (**Fig. S20**), a subtype previously found to be important in soils (37). Here, Form IE dominated the soil-derived Form I’s (1820 of 3876; **Fig. S3**) suggesting that it may have evolved to function in soil-like environments, especially those that lack substantial organic C inputs (e.g., deep soils, weathered rock, desert ecosystems (37,79).

In the absence of light, RuBisCO-mediated autotrophy is driven by chemical energy derived from the oxidation of electron donors (e.g., N, S, Fe, H_2_). Here, we found evidence of the oxidation of all four electron donors across all soil types, but abundance and expression differed across soil depth and ecosystems (**Fig. 2, Fig. 4**). Specifically, these genes were significantly more abundant in deep (**Fig. 2B**) and saturated soils (**Fig. S12, S13**). This enrichment in saturated soils may reflect a higher availability of reduced inorganic compounds such as H_2_, Fe^2+^, and reduced sulfur species. In non-saturated soils, the increase with depth may reflect increased reliance on CO_2_ fixation, less heterotrophic competitors for niche space, or lower oxygen availability. While the underlying redox drivers remain uncertain without direct oxygen measurements, our results align with findings by Frey et al. (2022) (80), who observed an overrepresentation of nitrification genes in deep forest soils. This pattern suggests that deep soil microbiomes are specifically adapted to the low oxygen conditions that favor these chemolithoautotrophic pathways.

Hydrogenases that mediate aerobic H_2_ oxidation were most abundant in weathering granite and deep saturated soils (**Fig. 2, Fig. S12, S13**). Hydrogen has been recognized as a prevalent and important electron donor in soils (81–83), even at extremely low concentrations (84,85). Specific high-affinity NiFe hydrogenases are widespread across bacteria and are able to scavenge H_2_ efficiently even at atmospheric trace levels (84,85). These enzymes co-occur with RuBisCO Form IE in the microbiome of a C-limited desert soil (37). Here, deep soils and weathered granite (saprolite) had more abundant hydrogenases and CBB-associated RuBisCO (**Fig. 2, Fig. S12, S13**), indicating potential elevated autotrophic metabolism fueled by H_2_. As there is transcriptomic evidence of active energy-permitting NiFe hydrogenases across nearly all soil types analyzed here (**Fig. 4**), hydrogen may be a key inorganic electron donor supporting deep soil autotrophy.

## Conclusion

Understanding the ecological context of soil autotrophy is crucial for interpreting its role in C cycling and storage. Here, we show that the prevalence, community composition, and activity of autotrophic microorganisms vary with environmental factors, such as soil saturation, depth, and ecosystem type (**Fig. 2, S10**). Our findings show that microbial autotrophy may be a key metabolism in soils with limited organic C inputs (e.g., deep soils, saprolite) and saturated ecosystems (rice paddies). Other sequencing and isotope-labeling studies (20,28,32,36,38,86) have shown that the contribution of autotrophy-derived C in soil microbial biomass might be significant, but is likely strongly modulated by soil type and biogeochemical gradients. Autotrophy appears to be an underappreciated but critical process in terrestrial ecosystems and, as climate change progresses, comprehensive understanding of the ecology of microbial autotrophs is essential for accurate modeling of soil C formation and cycling.

## Data availability

All data included here are either provided in the Supplementary Information, Supplemental Files, or are publicly available through previously published studies (detailed in Supplementary Data Sheet A). A subset of the metagenomic and metatranscriptomic datasets are derived from ongoing studies not yet fully published; therefore, to ensure transparency and reproducibility, the complete gene database used in this study has been made publicly available through Zenodo at https://doi.org/10.5281/zenodo.20329076.

## List of abbreviations

C: Carbon
SOM: soil organic matter
MAOM: mineral-associated organic matter
CBB: Calvin-Benson-Bassham
RuBisCO: ribulose-1,5-bisphosphate carboxylase/oxygenase
RLP: RuBisCO-like protein
WL: Wood-Ljungdahl
3HP: 3-hydroxypropionate
3HP/4HB: 3-Hydroxypropionate/4-hydroxybutyrate
rTCA: reverse tricarboxylic acid

## Contributions

ANK, JPR, and JFB conceptualized the study. ANK analyzed and curated the dataset, prepared all final figures and tables, and wrote the original draft. ALJ contributed to the marine RuBisCO data comparison analysis, and PIP, KEC, BCK, PMT, and MV contributed datasets and contributed to methodology. All authors reviewed the final manuscript.

## Declarations

### Ethics approval and consent to participate

Non-applicable.

### Consent for publication

Non-applicable.

## References

1. Friedlingstein P, O’Sullivan M, Jones MW, Andrew RM, Bakker DCE, Hauck J, et al. Global Carbon Budget 2025 [Internet]. ESSD – Anthroposphere/Energy and anthropogenic emissions; 2025 [cited 2026 Mar 20]. Available from: https://essd.copernicus.org/preprints/essd-2025-659/ doi:10.5194/essd-2025-659

2. Sanderman J, Hengl T, Fiske GJ. Soil carbon debt of 12,000 years of human land use. Proc Natl Acad Sci. 2018 Feb 13;115(7):E1700–E1700. doi:10.1073/pnas.1800925115

3. Paustian K, Lehmann J, Ogle S, Reay D, Robertson GP, Smith P. Climate-smart soils. Nature. 2016 Apr;532(7597):49–57. doi:10.1038/nature17174

4. Smith P, Martino D, Cai Z, Gwary D, Janzen H, Kumar P, et al. Greenhouse gas mitigation in agriculture. Philos Trans R Soc B Biol Sci. 2007 Sep 6;363(1492):789– 813. doi:10.1098/rstb.2007.2184

5. Fargione JE, Bassett S, Boucher T, Bridgham SD, Conant RT, Cook-Patton SC, et al. Natural climate solutions for the United States. Sci Adv. 2018 Nov 14;4(11):eaat1869. doi:10.1126/sciadv.aat1869

6. Domeignoz-Horta LA, Shinfuku M, Junier P, Poirier S, Verrecchia E, Sebag D, et al. Direct evidence for the role of microbial community composition in the formation of soil organic matter composition and persistence. ISME Commun. 2021 Dec 6;1(1):64. doi:10.1038/s43705-021-00071-7

7. Domeignoz-Horta LA, Pold G, Liu XJA, Frey SD, Melillo JM, DeAngelis KM. Microbial diversity drives carbon use efficiency in a model soil. Nat Commun. 2020 Dec 23;11(1):3684. doi:10.1038/s41467-020-17502-z PubMed PMID: 32703952.

8. Liang C, Amelung W, Lehmann J, Kästner M. Quantitative assessment of microbial necromass contribution to soil organic matter. Glob Change Biol. 2019;25(11):3578–90. doi:10.1111/gcb.14781

9. Kallenbach CM, Frey SD, Grandy AS. Direct evidence for microbial-derived soil organic matter formation and its ecophysiological controls. Nat Commun. 2016 Nov 28;7(1):13630. doi:10.1038/ncomms13630

10. Wang B, An S, Liang C, Liu Y, Kuzyakov Y. Microbial necromass as the source of soil organic carbon in global ecosystems. Soil Biol Biochem. 2021 Nov 1;162:108422. doi:10.1016/j.soilbio.2021.108422

11. Klink S, Keller AB, Wild AJ, Baumert VL, Gube M, Lehndorff E, et al. Stable isotopes reveal that fungal residues contribute more to mineral-associated organic matter pools than plant residues. Soil Biol Biochem. 2022 May;168(November 2021):108634. doi:10.1016/j.soilbio.2022.108634

12. Liang C, Schimel JP, Jastrow JD. The importance of anabolism in microbial control over soil carbon storage. Nat Microbiol. 2017 Aug 25;2(8):17105. doi:10.1038/nmicrobiol.2017.105 PubMed PMID: 28741607.

13. Cotrufo MF, Lavallee JM. Soil organic matter formation, persistence, and functioning: A synthesis of current understanding to inform its conservation and regeneration. In: Advances in Agronomy [Internet]. Elsevier; 2022 [cited 2024 Oct 22]. p. 1–66. Available from: https://linkinghub.elsevier.com/retrieve/pii/S0065211321001048 doi:10.1016/bs.agron.2021.11.002

14. Waring BG, Sulman BN, Reed S, Smith AP, Averill C, Creamer CA, et al. From pools to flow: The PROMISE framework for new insights on soil carbon cycling in a changing world. Glob Change Biol. 2020 Dec 16;26(12):6631–43. doi:10.1111/gcb.15365 PubMed PMID: 33064359.

15. Keller AB, Brzostek ER, Craig ME, Fisher JB, Phillips RP. Root-derived inputs are major contributors to soil carbon in temperate forests, but vary by mycorrhizal type. Ecol Lett. 2021;24(4):626–35. doi:10.1111/ele.13651

16. Whalen ED, Grandy AS, Sokol NW, Keiluweit M, Ernakovich J, Smith RG, et al. Clarifying the evidence for microbial-and plant-derived soil organic matter, and the path toward a more quantitative understanding. Glob Change Biol. 2022;28(24):7167–85. doi:10.1111/gcb.16413

17. Liao H, Hao X, Qin F, Delgado-Baquerizo M, Liu Y, Zhou J, et al. Microbial autotrophy explains large-scale soil CO fixation. Glob Change Biol. 2023;29(1):231–42. doi:10.1111/gcb.16452

18. Akinyede R, Taubert M, Schrumpf M, Trumbore S, Küsel K. Dark CO2 fixation in temperate beech and pine forest soils. Soil Biol Biochem. 2022 Feb;165:108526. doi:10.1016/j.soilbio.2021.108526

19. Xiao KQ, Ge TD, Wu XH, Peacock CL, Zhu ZK, Peng J, et al. Metagenomic and C tracing evidence for autotrophic microbial CO fixation in paddy soils. Environ Microbiol. 2021;23(2):924–33. doi:10.1111/1462-2920.15204

20. Yuan H, Ge T, Chen C, O’Donnell AG, Wu J. Significant Role for Microbial Autotrophy in the Sequestration of Soil Carbon. Appl Environ Microbiol. 2012 Apr;78(7):2328–36. doi:10.1128/AEM.06881-11

21. Prywes N, Phillips NR, Tuck OT, Valentin-Alvarado LE, Savage DF. Rubisco Function, Evolution, and Engineering. Annu Rev Biochem. 2023 Jun 20;92(Volume 92, 2023):385–410. doi:10.1146/annurev-biochem-040320-101244

22. Badger MR, Bek EJ. Multiple Rubisco forms in proteobacteria: their functional significance in relation to CO2 acquisition by the CBB cycle. J Exp Bot. 2008 May 1;59(7):1525–41. doi:10.1093/jxb/erm297

23. Banda DM, Pereira JH, Liu AK, Orr DJ, Hammel M, He C, et al. Novel bacterial clade reveals origin of form I Rubisco. Nat Plants. 2020 Sep;6(9):1158–66. doi:10.1038/s41477-020-00762-4

24. Kehl AJ, Taylor-Kearney L, Jaffe AL, Pereira JH, Lee J, Hammel M, et al. Diversity-driven biochemical survey reveals dimeric structural origin of rubisco [Internet]. bioRxiv; 2025 [cited 2025 Nov 20]. p. 2025.11.05.686826. Available from: https://www.biorxiv.org/content/10.1101/2025.11.05.686826v1 doi:10.1101/2025.11.05.686826

25. Selesi D, Schmid M, Hartmann A. Diversity of Green-Like and Red-Like Ribulose-1,5-Bisphosphate Carboxylase/Oxygenase Large-Subunit Genes (cbbL) in Differently Managed Agricultural Soils. Appl Environ Microbiol. 2005 Jan;71(1):175–84. doi:10.1128/AEM.71.1.175-184.2005

26. Tolli J, King GM. Diversity and Structure of Bacterial Chemolithotrophic Communities in Pine Forest and Agroecosystem Soils. Appl Environ Microbiol. 2005 Dec;71(12):8411–8. doi:10.1128/AEM.71.12.8411-8418.2005

27. Yuan H, Ge T, Zou S, Wu X, Liu S, Zhou P, et al. Effect of land use on the abundance and diversity of autotrophic bacteria as measured by ribulose-1,5-biphosphate carboxylase/oxygenase (RubisCO) large subunit gene abundance in soils. Biol Fertil Soils. 2013 Jul;49(5):609–16. doi:10.1007/s00374-012-0750-x

28. Xiao KQ, Bao P, Bao QL, Jia Y, Huang FY, Su JQ, et al. Quantitative analyses of ribulose-1,5-bisphosphate carboxylase/oxygenase (RubisCO) large-subunit genes (cbbL) in typical paddy soils. FEMS Microbiol Ecol. 2014 Jan 1;87(1):89–101. doi:10.1111/1574-6941.12193

29. Zhou ZF, Wei WL, Shi XJ, Liu YM, He XH, Wang MX. Twenty-six years of chemical fertilization decreased soil RubisCO activity and changed the ecological characteristics of soil cbbL-carrying bacteria in an entisol. Appl Soil Ecol. 2019 Sep;141:1–9. doi:10.1016/j.apsoil.2019.05.005

30. Song Y, Mei W, Li M, Wang X, Luo S, Feng Y, et al. Soil water content and RubisCO activity control the carbon storage in soil under different land uses in Sanjiang Plain, China. CATENA. 2024 Aug;243:108211. doi:10.1016/j.catena.2024.108211

31. Guo G, Kong W, Liu J, Zhao J, Du H, Zhang X, et al. Diversity and distribution of autotrophic microbial community along environmental gradients in grassland soils on the Tibetan Plateau. Appl Microbiol Biotechnol. 2015 Oct;99(20):8765–76. doi:10.1007/s00253-015-6723-x

32. Lynn TM, Ge T, Yuan H, Wei X, Wu X, Xiao K, et al. Soil Carbon-Fixation Rates and Associated Bacterial Diversity and Abundance in Three Natural Ecosystems. Microb Ecol. 2017 Apr 1;73(3):645–57. doi:10.1007/s00248-016-0890-x

33. Huang Q, Huang Y, Wang B, Dippold MA, Li H, Li N, et al. Metabolic pathways of CO2 fixing microorganisms determined C-fixation rates in grassland soils along the precipitation gradient. Soil Biol Biochem. 2022 Sep;172:108764. doi:10.1016/j.soilbio.2022.108764

34. Grüterich L, Woodhouse JN, Mueller P, Tiemann A, Ruscheweyh HJ, Sunagawa S, et al. Assessing environmental gradients in relation to dark CO2 fixation in estuarine wetland microbiomes. Appl Environ Microbiol. 2024 Dec 31;91(1):e02177–24. doi:10.1128/aem.02177-24

35. Yu H, He Z, Wang A, Xie J, Wu L, Van Nostrand JD, et al. Divergent Responses of Forest Soil Microbial Communities under Elevated CO2 in Different Depths of Upper Soil Layers. Appl Environ Microbiol. 2017 Dec 15;84(1):e01694–17. doi:10.1128/AEM.01694-17

36. Bu L, Peng Z, Tian J, Zhang X, Chen W, An D, et al. Core autotrophic microbes drive functional stability of soil cbbL-containing autotrophic microbes during desertification. Appl Soil Ecol. 2023 Oct;190:105027. doi:10.1016/j.apsoil.2023.105027

37. Ray AE, Zaugg J, Benaud N, Chelliah DS, Bay S, Wong HL, et al. Atmospheric chemosynthesis is phylogenetically and geographically widespread and contributes significantly to carbon fixation throughout cold deserts. ISME J. 2022 Nov 1;16(11):2547–60. doi:10.1038/s41396-022-01298-5

38. Bay SK, Waite DW, Dong X, Gillor O, Chown SL, Hugenholtz P, et al. Chemosynthetic and photosynthetic bacteria contribute differentially to primary production across a steep desert aridity gradient. ISME J. 2021 Nov;15(11):3339– 56. doi:10.1038/s41396-021-01001-0

39. Ji M, Greening C, Vanwonterghem I, Carere CR, Bay SK, Steen JA, et al. Atmospheric trace gases support primary production in Antarctic desert surface soil. Nature. 2017 Dec;552(7685):400–3. doi:10.1038/nature25014

40. Jaffe AL, Salcedo RSR, Dekas AE. Abundant and metabolically flexible bacterial lineages underlie a vast potential for rubisco-mediated carbon fixation in the dark ocean. Genome Biol. 2025 Jun 16;26(1):167. doi:10.1186/s13059-025-03625-3

41. Finn RD, Clements J, Eddy SR. HMMER web server: interactive sequence similarity searching. Nucleic Acids Res. 2011 Jul 1;39(suppl_2):W29–37. doi:10.1093/nar/gkr367

42. Boyd JA, Woodcroft BJ, Tyson GW. GraftM: a tool for scalable, phylogenetically informed classification of genes within metagenomes. Nucleic Acids Res. 2018 Jun 1;46(10):e59. doi:10.1093/nar/gky174

43. Rognes T, Flouri T, Nichols B, Quince C, Mahé F. VSEARCH: a versatile open source tool for metagenomics. PeerJ. 2016 Oct 18;4:e2584. doi:10.7717/peerj.2584

44. Katoh K. MAFFT: a novel method for rapid multiple sequence alignment based on fast Fourier transform. Nucleic Acids Res. 2002 Jul 15;30(14):3059–66. doi:10.1093/nar/gkf436 PubMed PMID: 12136088.

45. FastTree 2 – Approximately Maximum-Likelihood Trees for Large Alignments | PLOS ONE [Internet]. [cited 2024 Oct 23]. Available from: https://journals.plos.org/plosone/article?id=10.1371/journal.pone.0009490

46. Letunic I, Bork P. Interactive Tree of Life (iTOL) v6: recent updates to the phylogenetic tree display and annotation tool. Nucleic Acids Res. 2024 Jul 5;52(W1):W78–82. doi:10.1093/nar/gkae268

47. Steinegger M, Söding J. MMseqs2 enables sensitive protein sequence searching for the analysis of massive data sets. Nat Biotechnol. 2017 Nov 16;35(11):1026–8. doi:10.1038/nbt.3988 PubMed PMID: 29035372.

48. The UniProt Consortium. UniProt: a worldwide hub of protein knowledge. Nucleic Acids Res. 2019 Jan 8;47(D1):D506–15. doi:10.1093/nar/gky1049

49. Levy-Booth DJ, Hashimi A, Roccor R, Liu LY, Renneckar S, Eltis LD, et al. Genomics and metatranscriptomics of biogeochemical cycling and degradation of lignin-derived aromatic compounds in thermal swamp sediment. ISME J. 2020 Nov 2. doi:10.1038/s41396-020-00820-x

50. Garber AI, Nealson KH, Okamoto A, McAllister SM, Chan CS, Barco RA, et al. FeGenie: A Comprehensive Tool for the Identification of Iron Genes and Iron Gene Neighborhoods in Genome and Metagenome Assemblies. Front Microbiol. 2020 Jan 31;11. doi:10.3389/fmicb.2020.00037

51. Søndergaard D, Pedersen CNS, Greening C. HydDB: A web tool for hydrogenase classification and analysis. Sci Rep. 2016 Sep 27;6(1):34212. doi:10.1038/srep34212

52. Singleton CM, McCalley CK, Woodcroft BJ, Boyd JA, Evans PN, Hodgkins SB, et al. Methanotrophy across a natural permafrost thaw environment. ISME J. 2018 Oct;12(10):10. doi:10.1038/s41396-018-0065-5

53. Müller AL, Kjeldsen KU, Rattei T, Pester M, Loy A. Phylogenetic and environmental diversity of DsrAB-type dissimilatory (bi)sulfite reductases. ISME J. 2015 May 1;9(5):1152–65. doi:10.1038/ismej.2014.208

54. Lappan R, Shelley G, Islam ZF, Leung PM, Lockwood S, Nauer PA, et al. Molecular hydrogen in seawater supports growth of diverse marine bacteria. Nat Microbiol. 2023 Apr;8(4):581–95. doi:10.1038/s41564-023-01322-0

55. Buchfink B, Xie C, Huson DH. Fast and sensitive protein alignment using DIAMOND. Nat Methods. 2015 Jan;12(1):59–60. doi:10.1038/nmeth.3176

56. Langmead B, Salzberg SL. Fast gapped-read alignment with Bowtie 2. Nat Methods. 2012 Apr 4;9(4):357–9. doi:10.1038/nmeth.1923 PubMed PMID: 22388286.

57. Aroney STN, Newell RJP, Nissen JN, Camargo AP, Tyson GW, Woodcroft BJ. CoverM: Read alignment statistics for metagenomics [Internet]. arXiv; 2025 [cited 2025 Mar 25]. Available from: http://arxiv.org/abs/2501.11217 doi:10.48550/arXiv.2501.11217

58. Liao Y, Smyth GK, Shi W. featureCounts: an efficient general purpose program for assigning sequence reads to genomic features. Bioinformatics. 2014 Apr 1;30(7):923–30. doi:10.1093/bioinformatics/btt656

59. Team RC. R: A language and environment for statistical computing. R Found Stat Comput Vienna Austria [Internet]. 2021. Available from: https://www.r-project.org/

60. Kassambra A. ggpubr: “ggplot2” Based Publication Ready Plots [Internet]. 2023. Available from: https://CRAN.R-project.org/package=ggpubr

61. Oksanen J, Blanchet FG, Friendly M, Kindt R, Legendre P, Mcglinn D, et al. Package “vegan.” In. Dordrecht: Springer Netherlands; 2013. Available from: http://link.springer.com/10.1007/978-94-024-1179-9_301576 doi:10.1007/978-94-024-1179-9_301576

62. McMurdie PJ, Holmes S. phyloseq: An R Package for Reproducible Interactive Analysis and Graphics of Microbiome Census Data. Watson M, editor. PLoS ONE. 2013 Apr 22;8(4):e61217. doi:10.1371/journal.pone.0061217 PubMed PMID: 23630581.

63. Anderson MJ. A new method for non-parametric multivariate analysis of variance.

64. Wickham H. ggplot2: Elegant Graphics for Data Analysis. Springer-Verlag, New York; 2016.

65. Sharrar AM, Crits-Christoph A, Méheust R, Diamond S, Starr EP, Banfield JF. Bacterial Secondary Metabolite Biosynthetic Potential in Soil Varies with Phylum, Depth, and Vegetation Type. mBio. 2020 Jun 16;11(3):10.1128/mbio.00416-20. doi:10.1128/mbio.00416-20

66. Diamond S, Andeer PF, Li Z, Crits-Christoph A, Burstein D, Anantharaman K, et al. Mediterranean grassland soil C–N compound turnover is dependent on rainfall and depth, and is mediated by genomically divergent microorganisms. Nat Microbiol. 2019 Aug;4(8):1356–67. doi:10.1038/s41564-019-0449-y

67. Nicolas AM, Sieradzki ET, Pett-Ridge J, Banfield JF, Taga ME, Firestone MK, et al. A subset of viruses thrives following microbial resuscitation during rewetting of a seasonally dry California grassland soil. Nat Commun. 2023 Sep 20;14(1):5835. doi:10.1038/s41467-023-40835-4

68. Penev PI, Estera-Molina K, Allen GM, Sachdeva R, Lei S, Law KK, et al. The active subset of grassland soil microbiomes changes with soil depth, water availability and prominently features predatory bacteria and episymbionts [Internet]. bioRxiv; 2024 [cited 2025 Mar 25]. p. 2024.12.19.629468. Available from: https://www.biorxiv.org/content/10.1101/2024.12.19.629468v1 doi:10.1101/2024.12.19.629468

69. Lavy A, McGrath DG, Matheus Carnevali PB, Wan J, Dong W, Tokunaga TK, et al. Microbial communities across a hillslope-riparian transect shaped by proximity to the stream, groundwater table, and weathered bedrock. Ecol Evol. 2019;9(12):6869– 900. doi:10.1002/ece3.5254

70. Matheus Carnevali PB, Lavy A, Thomas AD, Crits-Christoph A, Diamond S, Méheust R, et al. Meanders as a scaling motif for understanding of floodplain soil microbiome and biogeochemical potential at the watershed scale. Microbiome. 2021 May 22;9(1):121. doi:10.1186/s40168-020-00957-z

71. Sorensen PO, Beller HR, Bill M, Bouskill NJ, Hubbard SS, Karaoz U, et al. The Snowmelt Niche Differentiates Three Microbial Life Strategies That Influence Soil Nitrogen Availability During and After Winter. Front Microbiol. 2020 May 15;11. doi:10.3389/fmicb.2020.00871

72. Crits-Christoph A, Diamond S, Al-Shayeb B, Valentin-Alvarado L, Banfield JF. A widely distributed genus of soil Acidobacteria genomically enriched in biosynthetic gene clusters. ISME Commun. 2022 Dec 1;2(1):70. doi:10.1038/s43705-022-00140-5

73. Valentin-Alvarado LE, Appler KE, De Anda V, Schoelmerich MC, West-Roberts J, Kivenson V, et al. Asgard archaea modulate potential methanogenesis substrates in wetland soil. Nat Commun. 2024 Jul 31;15(1):6384. doi:10.1038/s41467-024-49872-z

74. Valentin-Alvarado LE, Shi LD, Appler KE, Crits-Christoph A, Anda VD, Adler BA, et al. Complete genomes of Asgard archaea reveal diverse integrated and mobile genetic elements. Genome Res. 2024 Oct 1;34(10):1595–609. doi:10.1101/gr.279480.124 PubMed PMID: 39406503.

75. Voutsinos MY, West-Roberts JA, Sachdeva R, Moreau JW, Banfield JF. Weathered granites and soils harbour microbes with lanthanide-dependent methylotrophic enzymes. BMC Biol. 2024 Feb 19;22(1):41. doi:10.1186/s12915-024-01841-0

76. Kehl AJ, Taylor-Kearney L, Jaffe AL, Pereira JH, Lee J, Hammel M, et al. Diversity-driven biochemical survey reveals dimeric structural origin of rubisco [Internet]. bioRxiv; 2025 [cited 2026 Jan 8]. p. 2025.11.05.686826. Available from: https://www.biorxiv.org/content/10.1101/2025.11.05.686826v1 doi:10.1101/2025.11.05.686826

77. Brewer TE, Aronson EL, Arogyaswamy K, Billings SA, Botthoff JK, Campbell AN, et al. Ecological and Genomic Attributes of Novel Bacterial Taxa That Thrive in Subsurface Soil Horizons. mBio. 2019 Oct;10(5):10.1128/mbio.01318-19. doi:10.1128/mbio.01318-19

78. Fierer N. Embracing the unknown: Disentangling the complexities of the soil microbiome. Nat Rev Microbiol. 2017;15(10):579–90. doi:10.1038/nrmicro.2017.87

79. Harrison K, Rapp JZ, Jaffe AL, Deming JW, Young J. Chemoautotrophy in subzero environments and the potential for cold-adapted Rubisco. Lotti M, editor. Appl Environ Microbiol. 2025 Jun 18;91(6):e00604–25. doi:10.1128/aem.00604-25

80. Frey B, Varliero G, Qi W, Stierli B, Walthert L, Brunner I. Shotgun Metagenomics of Deep Forest Soil Layers Show Evidence of Altered Microbial Genetic Potential for Biogeochemical Cycling. Front Microbiol. 2022 Mar 1;13. doi:10.3389/fmicb.2022.828977

81. Piché-Choquette S, Constant P. Molecular Hydrogen, a Neglected Key Driver of Soil Biogeochemical Processes. Appl Environ Microbiol. 2019 Mar 6;85(6):e02418–18. doi:10.1128/AEM.02418-18

82. Jordaan K, Lappan R, Dong X, Aitkenhead IJ, Bay SK, Chiri E, et al. Hydrogen-Oxidizing Bacteria Are Abundant in Desert Soils and Strongly Stimulated by Hydration. mSystems. 2020 Nov 17;5(6):10.1128/msystems.01131-20. doi:10.1128/msystems.01131-20

83. Greening C, Constant P, Hards K, Morales SE, Oakeshott JG, Russell RJ, et al. Atmospheric Hydrogen Scavenging: from Enzymes to Ecosystems. Appl Environ Microbiol. 2015 Feb 15;81(4):1190–9. doi:10.1128/AEM.03364-14

84. Greening C, Berney M, Hards K, Cook GM, Conrad R. A soil actinobacterium scavenges atmospheric H2 using two membrane-associated, oxygen-dependent [NiFe] hydrogenases. Proc Natl Acad Sci. 2014 Mar 18;111(11):4257–61. doi:10.1073/pnas.1320586111

85. Greening C, Carere CR, Rushton-Green R, Harold LK, Hards K, Taylor MC, et al. Persistence of the dominant soil phylum Acidobacteria by trace gas scavenging. Proc Natl Acad Sci. 2015 Aug 18;112(33):10497–502. doi:10.1073/pnas.1508385112

86. Berg IA. Ecological Aspects of the Distribution of Different Autotrophic CO2 Fixation Pathways. Appl Environ Microbiol. 2011 Mar 15;77(6):1925–36. doi:10.1128/AEM.02473-10

