## Supplementary Information for "Microbial autotrophy is widespread across soils and most prevalent in deep and saturated environments"

#### This PDF file includes:

| | |
|---|---|
| Supplementary Text | 2-3 |
| Supplementary Figures 1-21 | 4-24 |
| Supplementary Tables 1-3 | 25-26 |
| Supplementary Dataset Information | 27 |
| References | 28 |

#### Other supporting materials for this manuscript include the following:

Supplementary Data File 1
Supplementary Data File 2
Supplementary Data File 3

### Supplementary text

*Ensuring metatranscriptomic mapping methods did not result in spurious results*

For this work, we chose to map metatranscriptomic reads to the entire gene dataset, instead of mapping to only genes recovered from metagenomic reads recovered from the same sites represented in the associated metatranscriptome. To ensure that this did not result in mismapping or spurious analyses, we first analysed the number of metatranscriptomic reads from a metatranscriptome that map to genes recovered from the same field site or project and found that most transcripts were indeed mapping to genes recovered from the same field site (**Table S3**). Metatranscriptomes from the Hopland field site (California grassland) had the lowest % of transcripts mapping to genes from the same field site, but most transcripts mapping to genes from other field sites were mapping to another grassland field site not far from Hopland (Angelo). Further, most East River transcripts mapping to genes recovered from a different field site were mapping to genes from other non-saturated soils. Saturated ecosystem (rice paddies and vernal pools) metatranscriptomes consistently mapped to genes recovered from the same field site. Further, we also did one test analysis by mapping vernal pool metatranscriptomes to only genes recovered from the vernal pools to ensure we were seeing the same broad trends in gene transcript abundance, and we found that trends were broadly similar. For example, in this test, the highest activity of *prk*, RuBisCO Form I, and *foxY* was in mid-depth saturated soils, and transcripts mapped to *foxZ* in shallow and mid-depth saturated soils but were absent in deep saturated soils (**Fig. S21**). This check confirmed that our metatranscriptomic mapping techniques here did not lead to spurious results.

*Diversity of Form IV RuBisCO in soils (RLPs)*

In this meta-analysis, we found substantial compositional (**Fig. S7-9**) and sequence diversity in RuBisCO encoded in the soil microbiome (**Fig. 1**) and across terrestrial and marine ecosystems (**Fig. 5**). Form IV RuBisCO exhibited a large amount of taxonomic diversity and the highest phylogenetic diversity among RuBisCO (**Fig. 1**). These Form IV RuBisCO proteins, also known as RuBisCO-like proteins (RLPs), are not involved in the CBB cycle but have been linked to a variety of organism-specific functions, including roles in S metabolism such as methionine salvage and thiosulfate oxidation (1). The function of RLPs in some organisms, though, remains unknown (2). Our findings show that RLPs may be ecosystem-dependent, with distinct Form IV clades more commonly represented in either soil or marine metagenomes (**Fig. 5**). This pattern suggests that RLPs may have specialized roles that drive environmental variation in the abundance of distinct RLP clades. The functional diversity and ecosystem specificity of RLPs could have important implications for S cycling and broader ecosystem processes, underscoring the need for further study to clarify their ecological roles.

### Supplementary figures

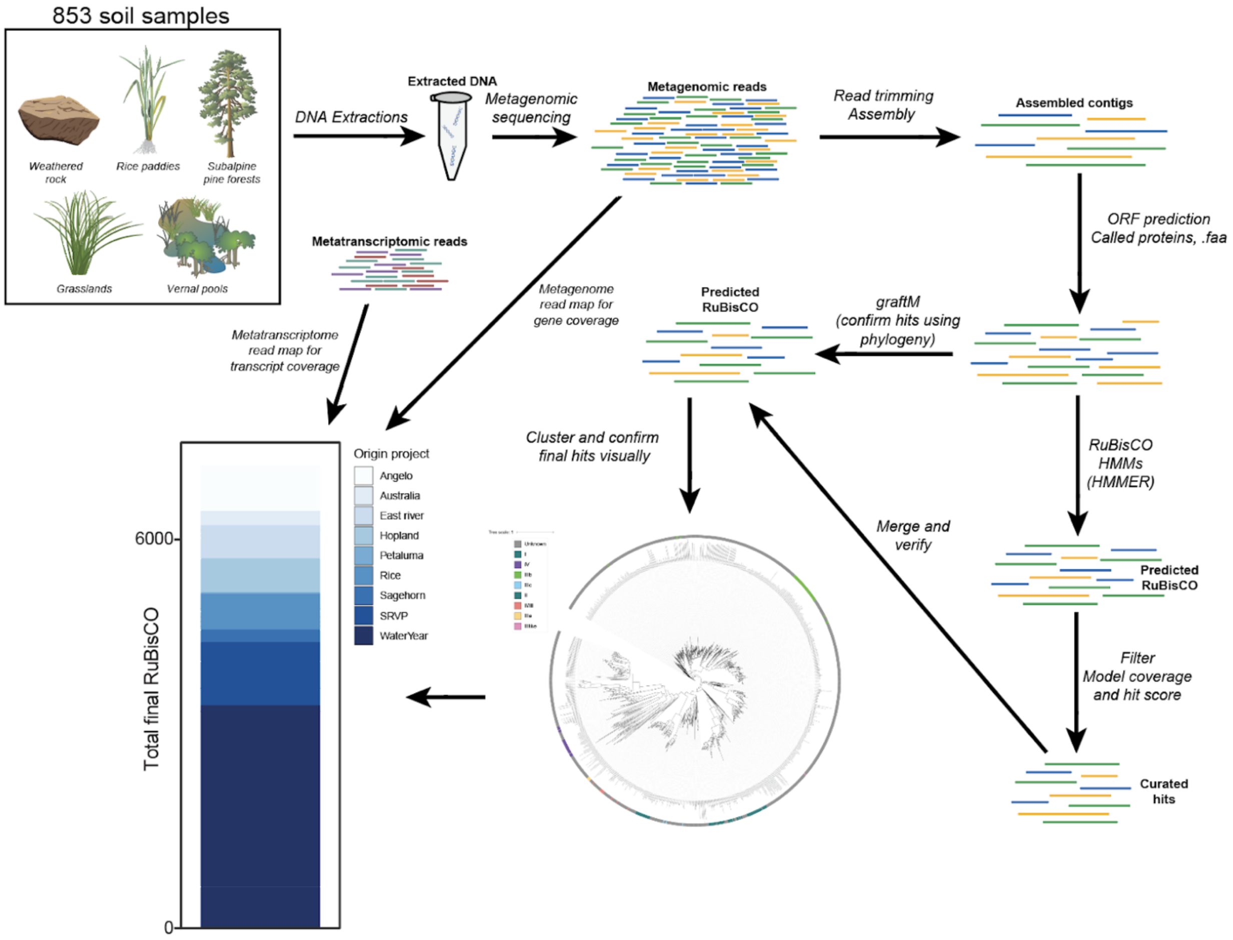


**Fig. S1.** Overview of bioinformatic pipeline for recovering RuBisCO sequences from metagenomic dataset. Genes were called from protein sequences and HMMs were run on all proteins for preliminary annotations, which were filtered to those with at least 50% HMM coverage and a score >100. Form-level annotations were confirmed using graftM and any final conflicting form-level annotations were validated using phylogenetic tree validation. The final RuBisCO protein set was filtered to those ≥400 amino acids long, resulting in 7140 final RuBisCO proteins used for analyses here.

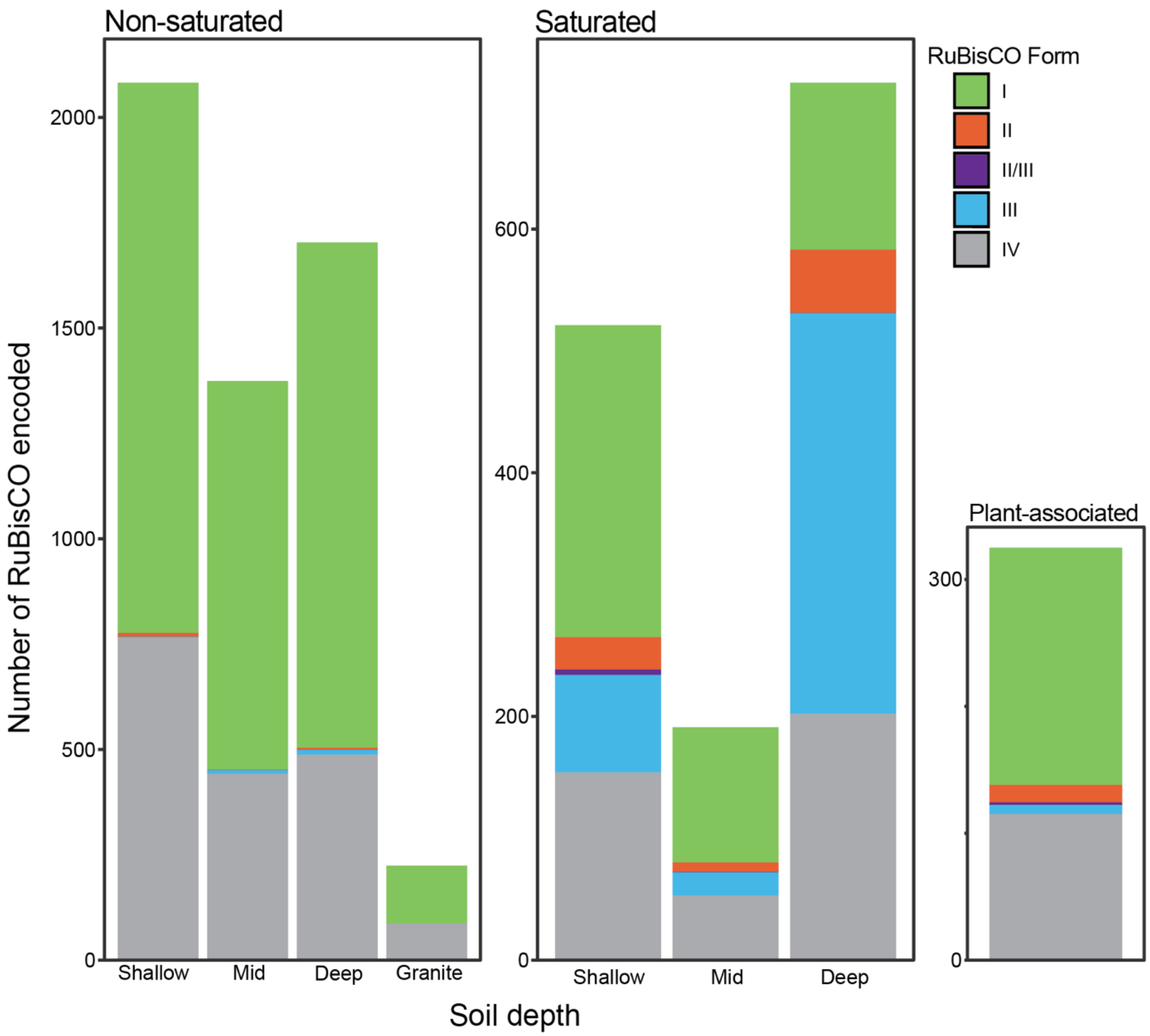


**Fig. S2.** Number of unique RuBisCO in final soil dataset from across ecosystem types (non-saturated, saturated, plant-associated) and soil depth (shallow = ≤25 cm, mid = 25-50 cm, deep = >50 cm, granite indicates a weathering granite profile, plant-associated samples not separated by root depth).

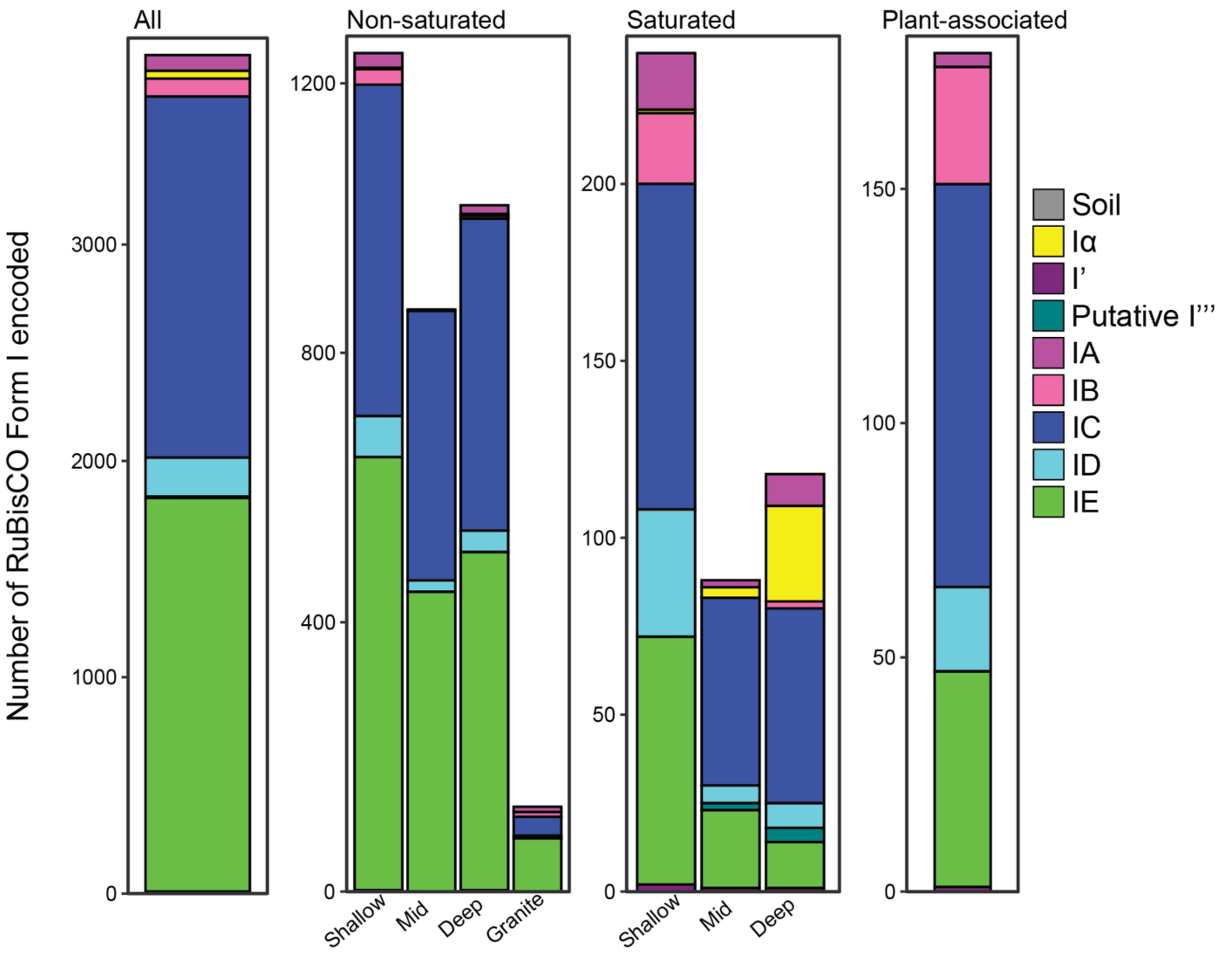


**Fig. S3.** Phylogenetically-verified, Form I annotations of Form I's in the soil-derived dataset, separated by metagenome soil origin ecosystem type (permanently saturated vs. non-saturated ecosystems) and soil depth (shallow = ≤25 cm, mid = 25-50 cm, deep = >50 cm).

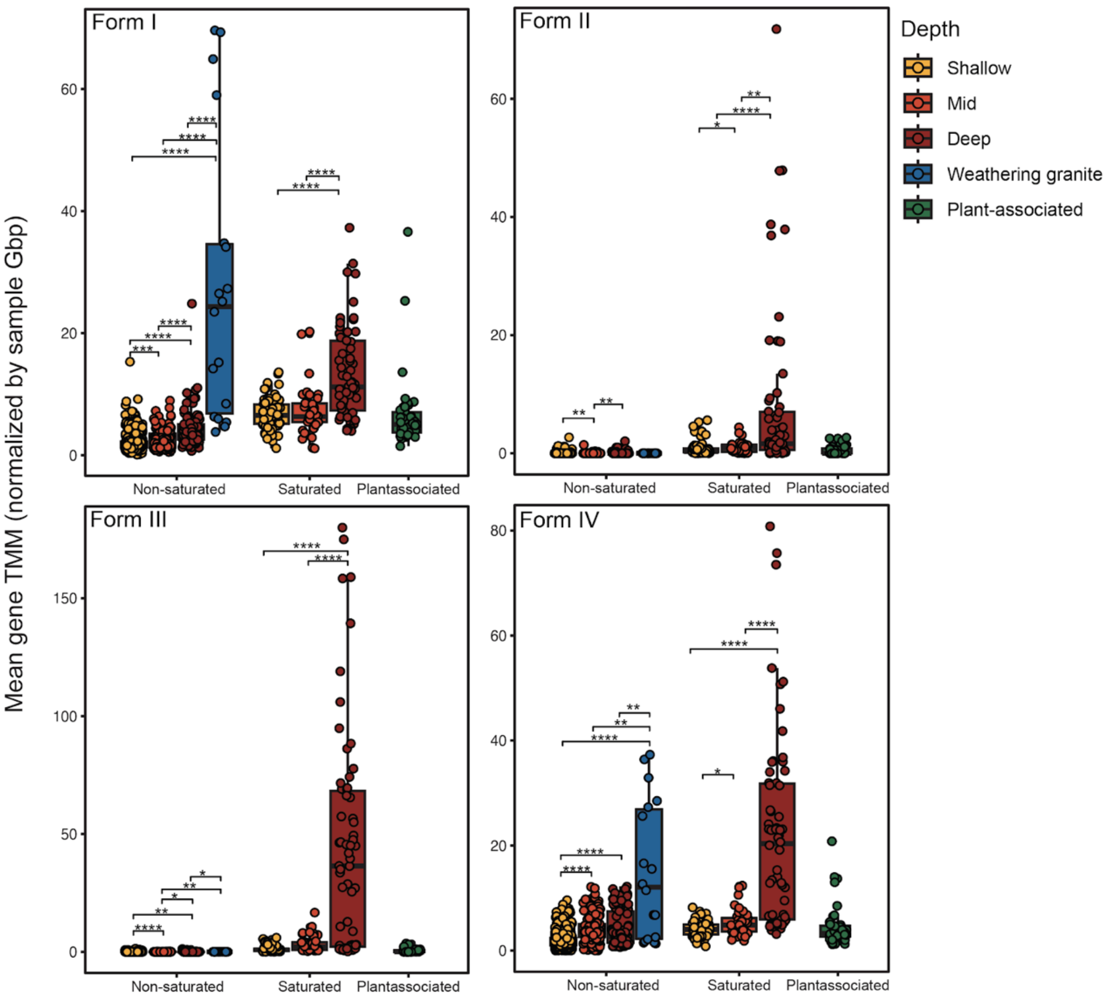


**Fig. S4.** Gene TMM of different forms of RuBisCO across samples. Points represent individual soil samples. The lower and upper hinges of the boxplots represent the 25th and 75th percentiles, respectively, and the middle line is the median. The whiskers extend from the median by 1.5x the interquartile range. Points represent individual samples. Significant differences between soil depths indicated with asterisks as indicated by Wilcoxon rank-sum test. *p<0.05, **p<0.01, ***p<0.001, ****p<0.0001.

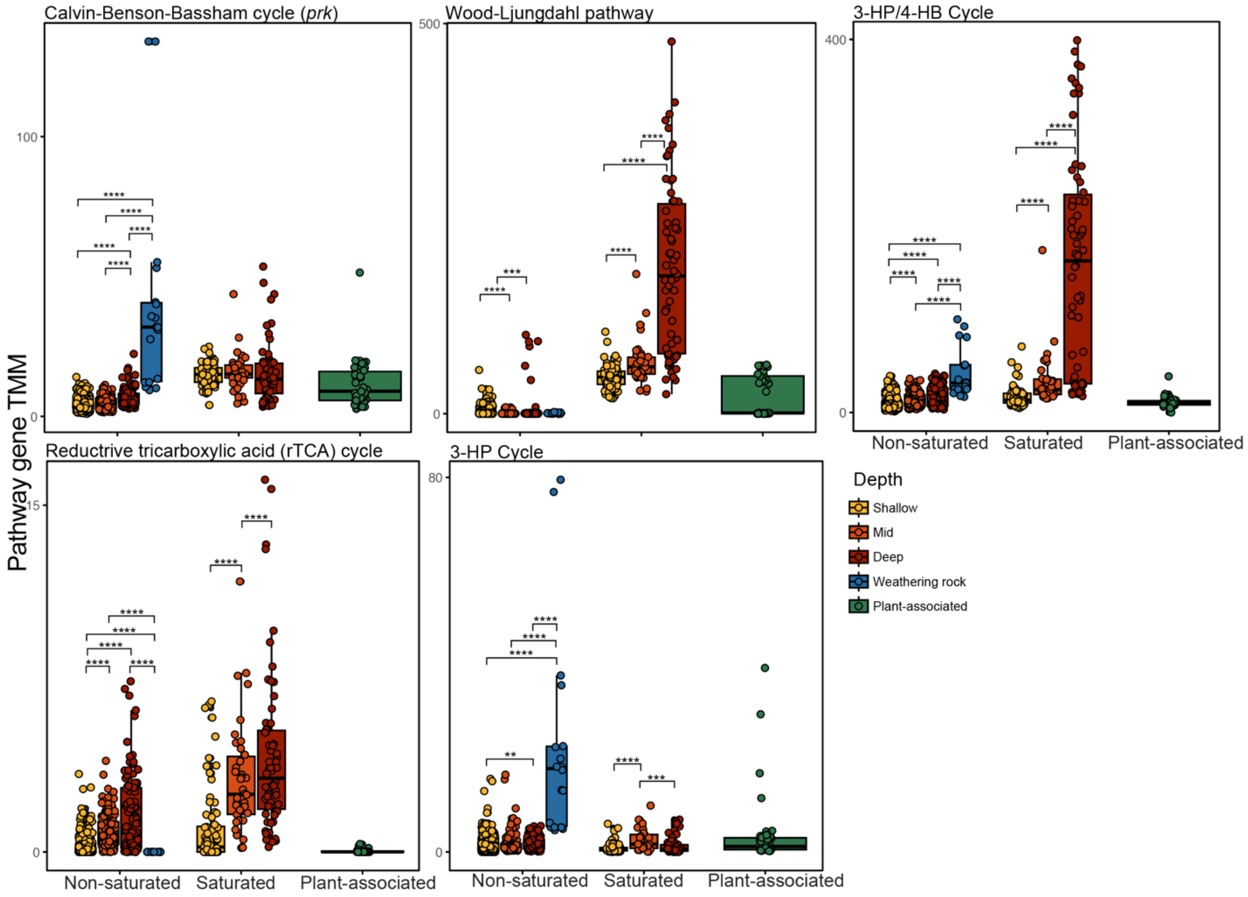


**Fig. S5.** Gene TMM of different forms of genes of different autotrophic pathways across samples. Points represent individual soil samples. The lower and upper hinges of the boxplots represent the 25th and 75th percentiles, respectively, and the middle line is the median. The whiskers extend from the median by 1.5x the interquartile range. Points represent individual samples. Significant differences between soil depths indicated with asterisks as indicated by Wilcoxon rank-sum test. *p<0.05, **p<0.01, ***p<0.001, ****p<0.0001.

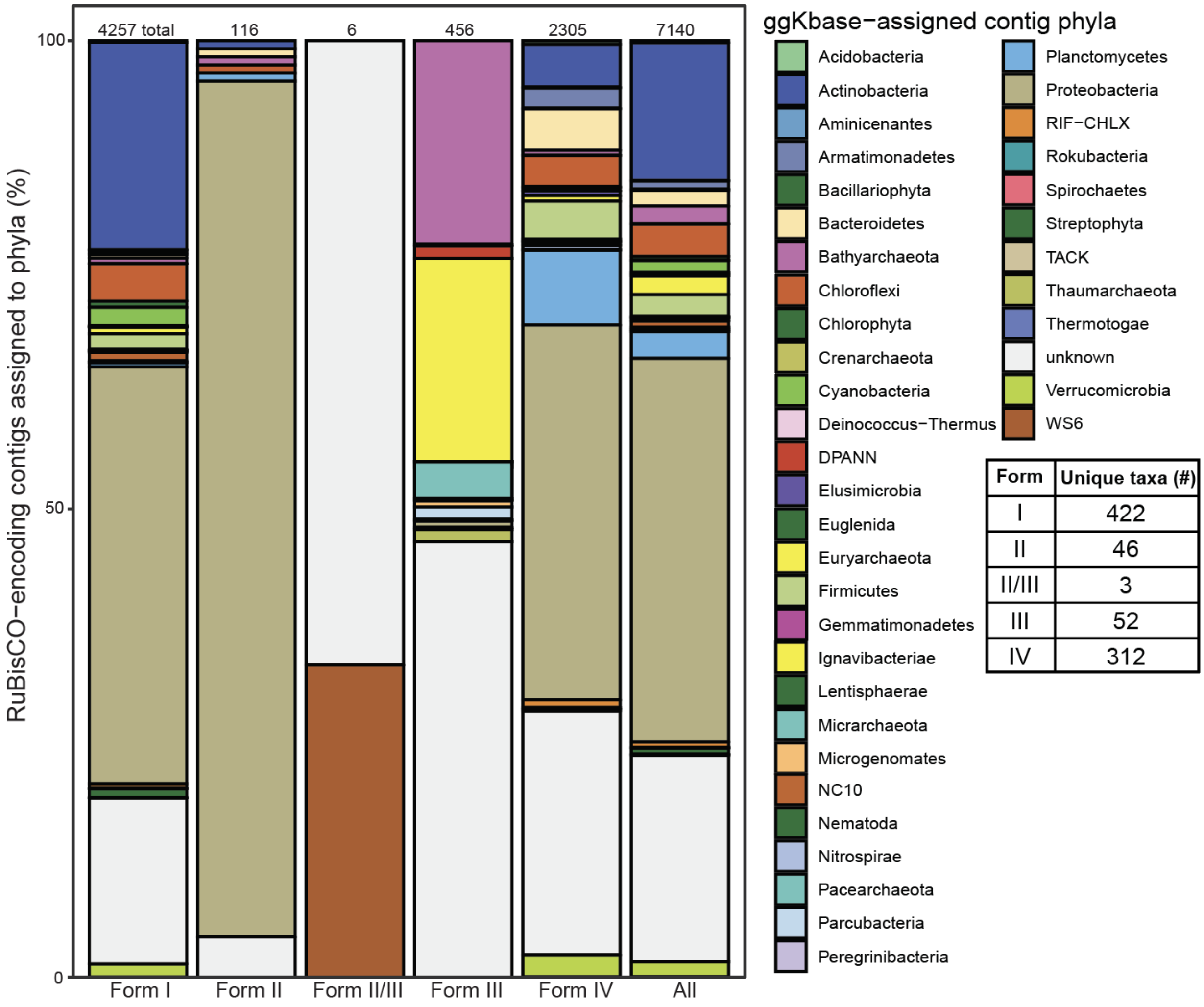


| Form | Unique taxa (#) |
|---|---|
| I | 422 |
| II | 46 |
| II/III | 3 |
| III | 52 |
| IV | 312 |

**Fig. S6.** Phyla-level taxonomy of RuBisCO sequences across all RuBisCO forms. Table includes number of unique lineages encoding each form.

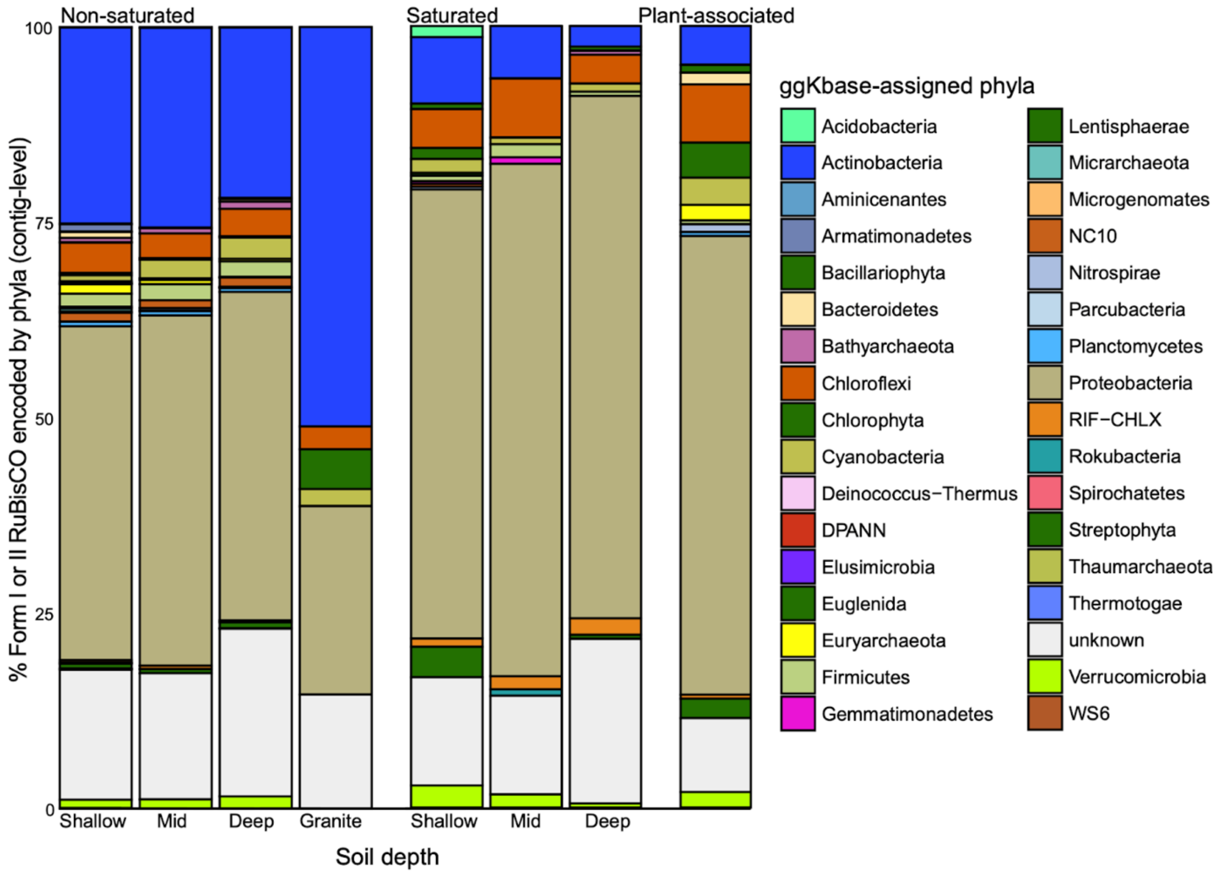


**Fig. S7.** Phyla-level taxonomy of CBB-associated RuBisCO Form I and II sequences encoded across ecosystem type (non-saturated, saturated, plant-associated) and depth (shallow = ≤25 cm, mid = 25-50 cm, deep = >50 cm, granite indicates a weathering granite profile, plant-associated samples not separated by root depth).

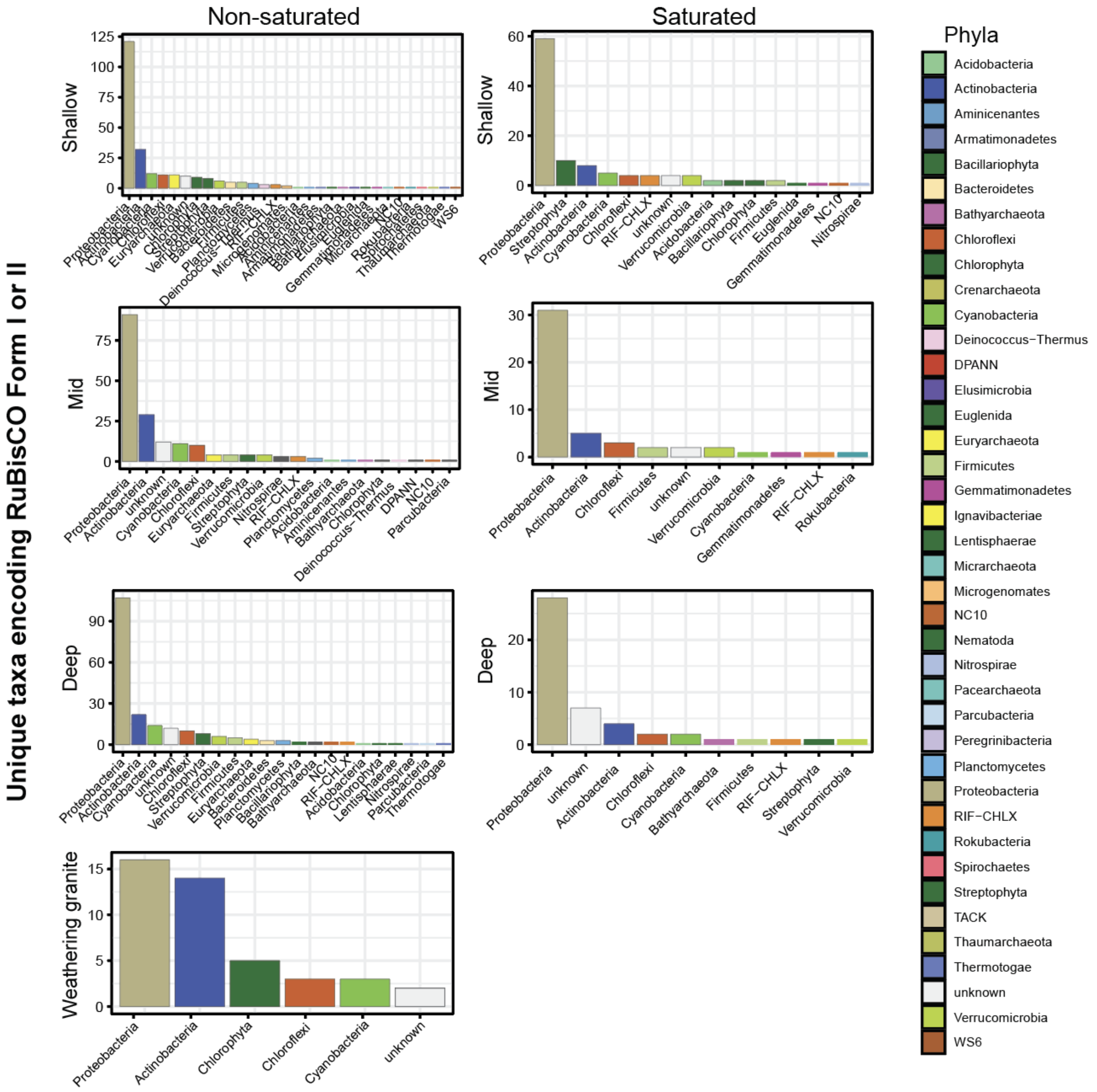


**Fig. S8.** Number of unique taxa within each phyla that encode RuBisCO Form I or II across all soil conditions and depths.

A

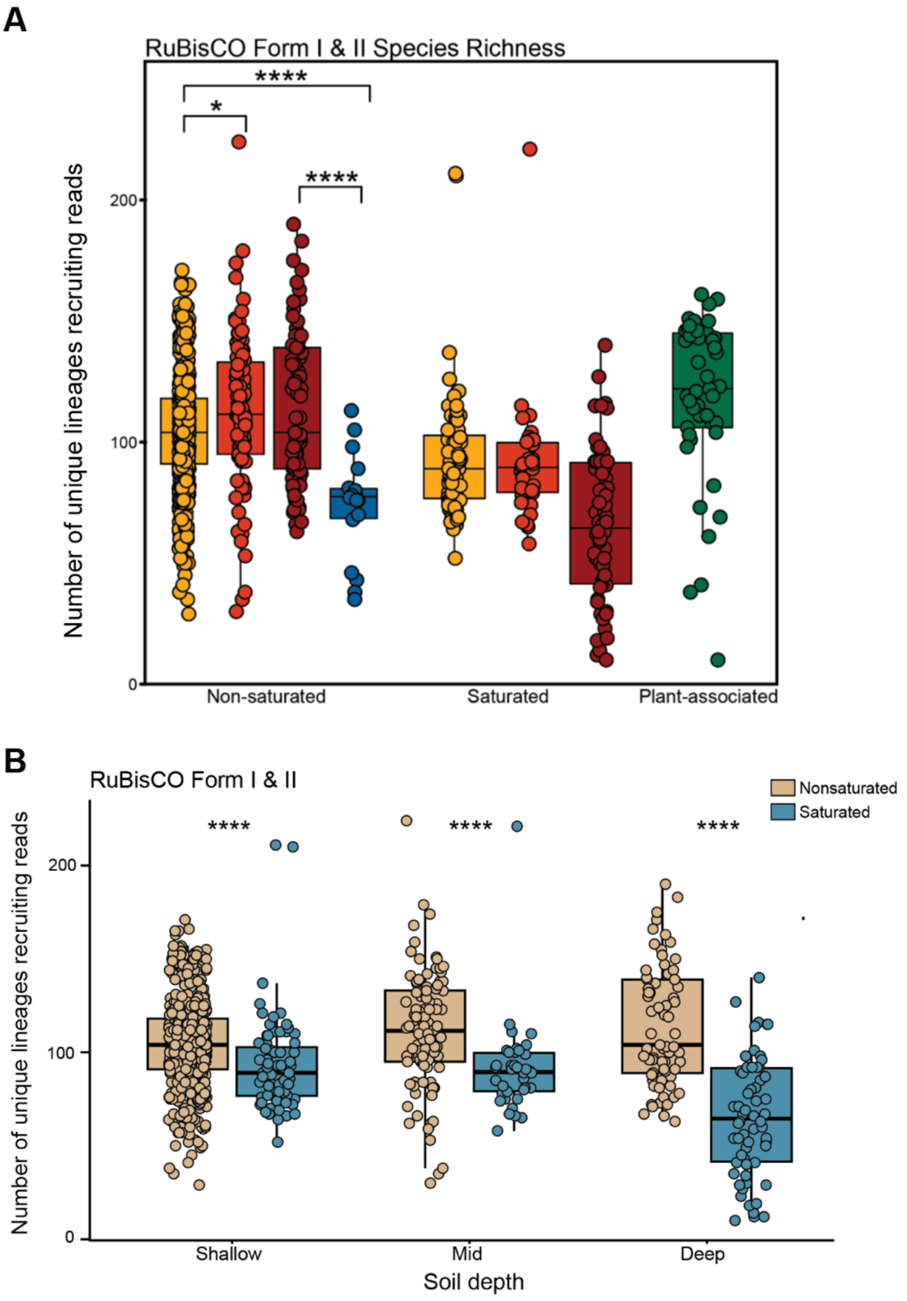


**Fig. S9.** Number of unique microbial lineages encoding RuBisCO Form I & II that recruit metagenomic reads (via read mapping) across soil types and depth. The lower and upper hinges of the boxplots represent the 25th and 75th percentiles, respectively, and the middle line is the median. The whiskers extend from the median by 1.5x the interquartile range. Points represent individual samples. Significant differences between conditions indicated with asterisks as indicated by Wilcoxon rank-sum test. *p<0.05, **p<0.01, ***p<0.001, ****p<0.0001.

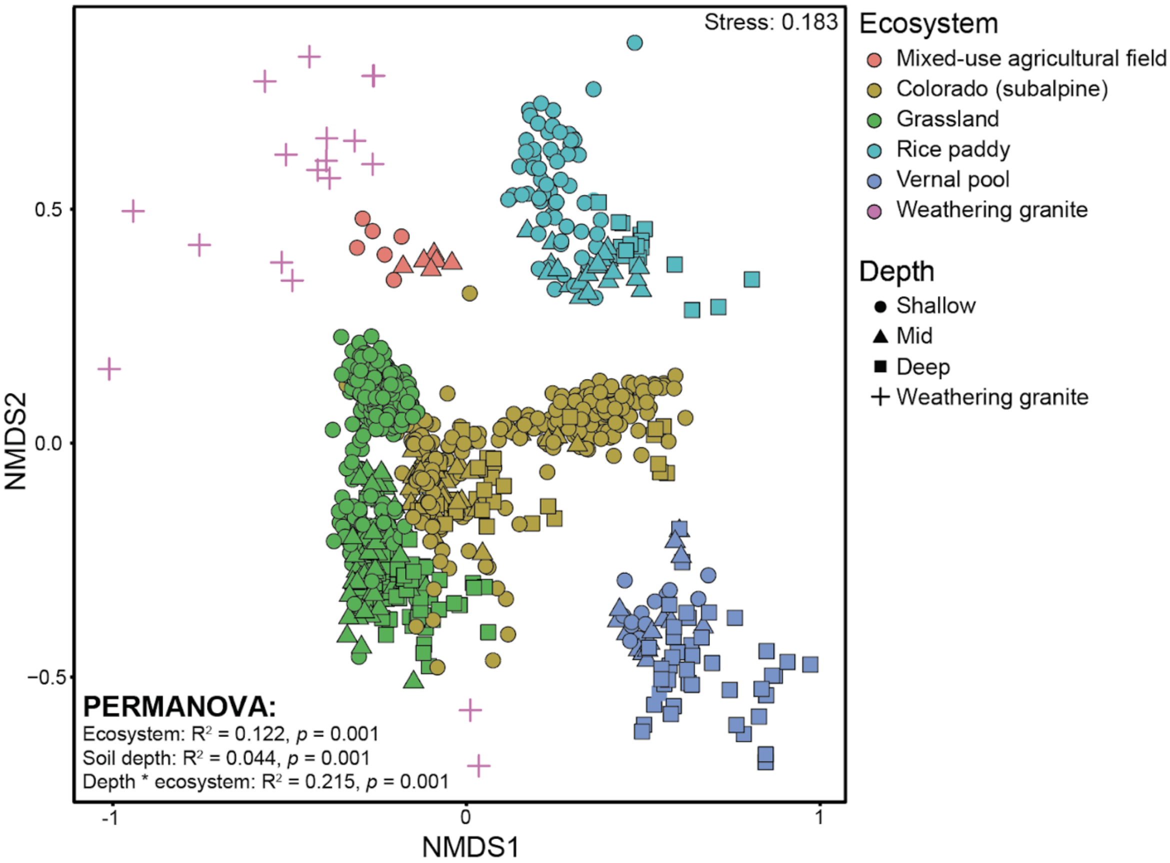


**Fig. S10.** Non-metric multidimensional scaling (NMDS) ordination of Bray-Curtis dissimilarity between soil samples based on normalized autotrophy gene abundance profiles (including all pathways), including all bulk soil samples.

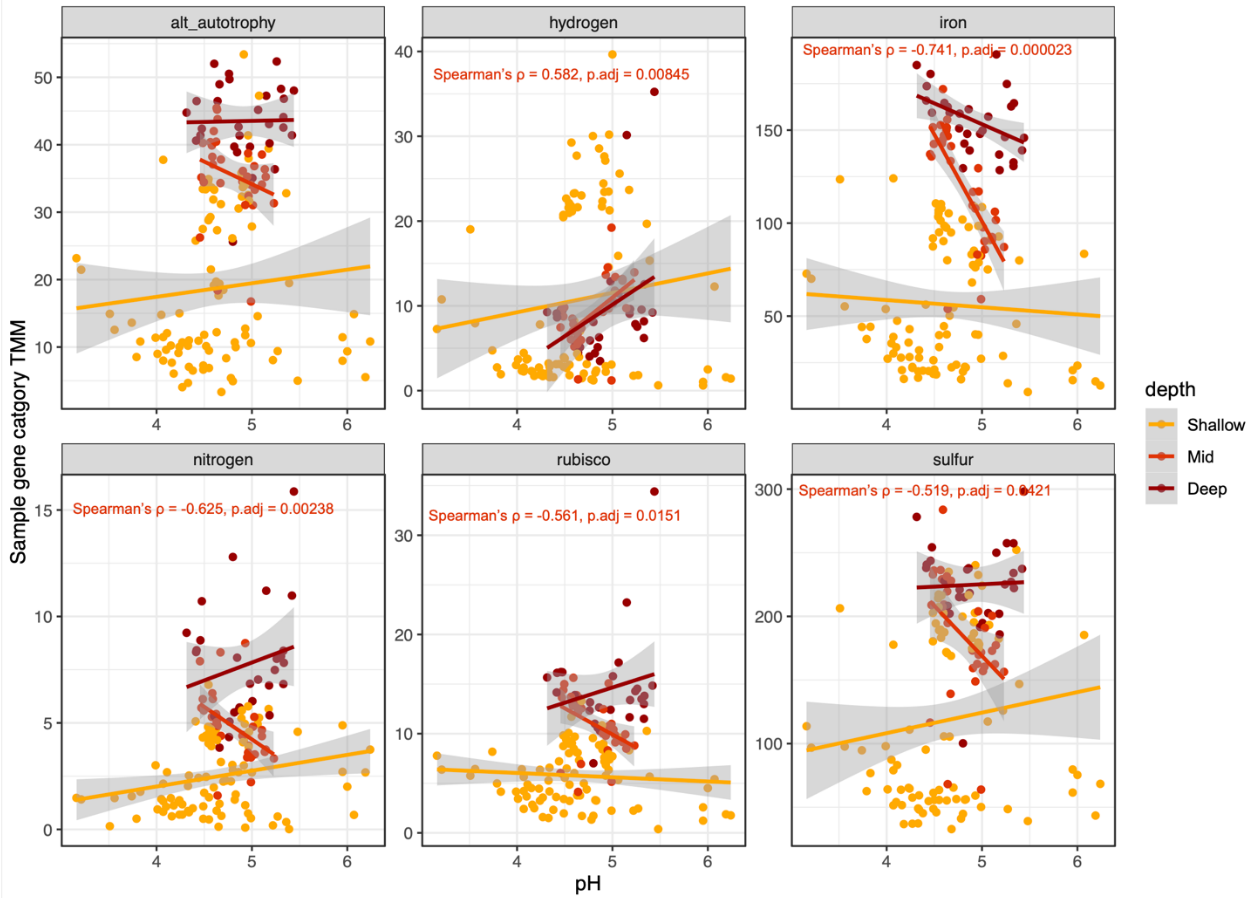


**Fig. S11.** Given category gene abundance (e.g., alternate autotrophy pathways, rubisco, sulfur oxidation) plotted against reported soil pH. Spearman's rho values are reported with Bonferroni-adjusted p-values, colored by soil depth, if significant. Shaded area shoes 95% confidence interval of linear model.

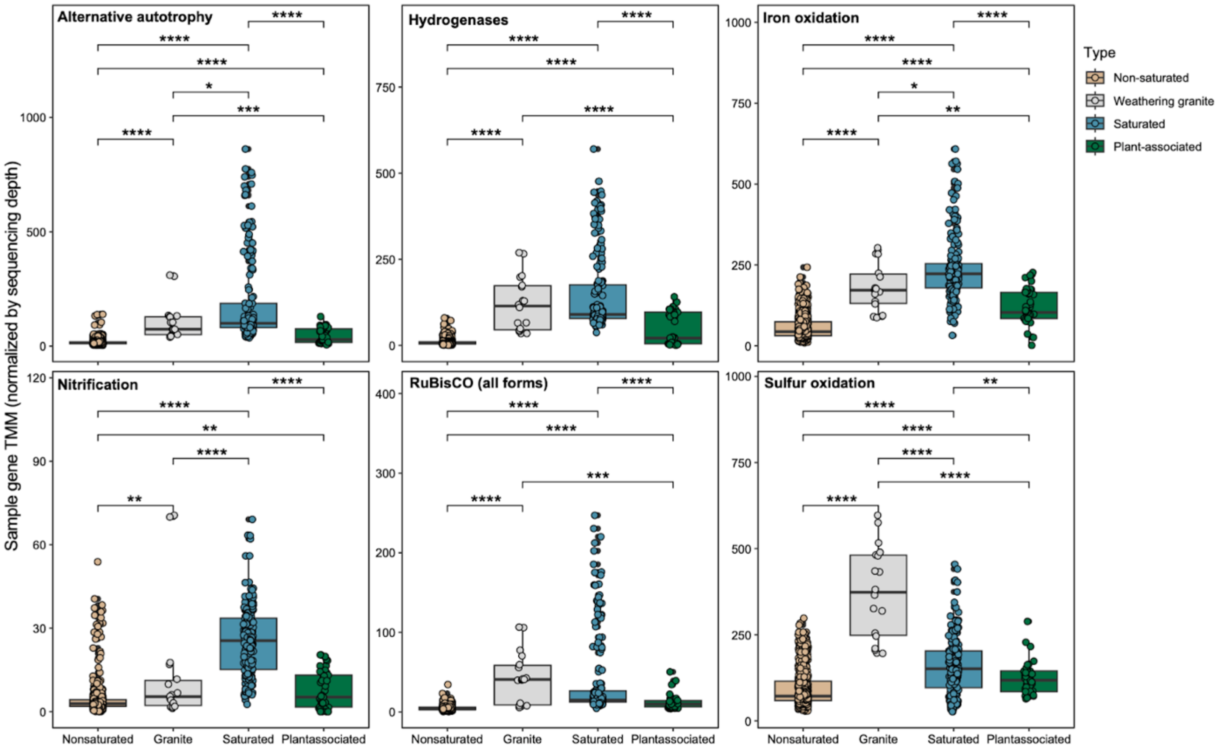


**Fig. S12.** Summed gene TMM of different metabolic or gene categories across samples. Points represent individual soil samples. The lower and upper hinges of the boxplots represent the 25th and 75th percentiles, respectively, and the middle line is the median. The whiskers extend from the median by 1.5x the interquartile range. Points represent individual samples. Significant differences between soil depths indicated with asterisks as indicated by Wilcoxon rank-sum test. *p<0.05, **p<0.01, ***p<0.001, ****p<0.0001.

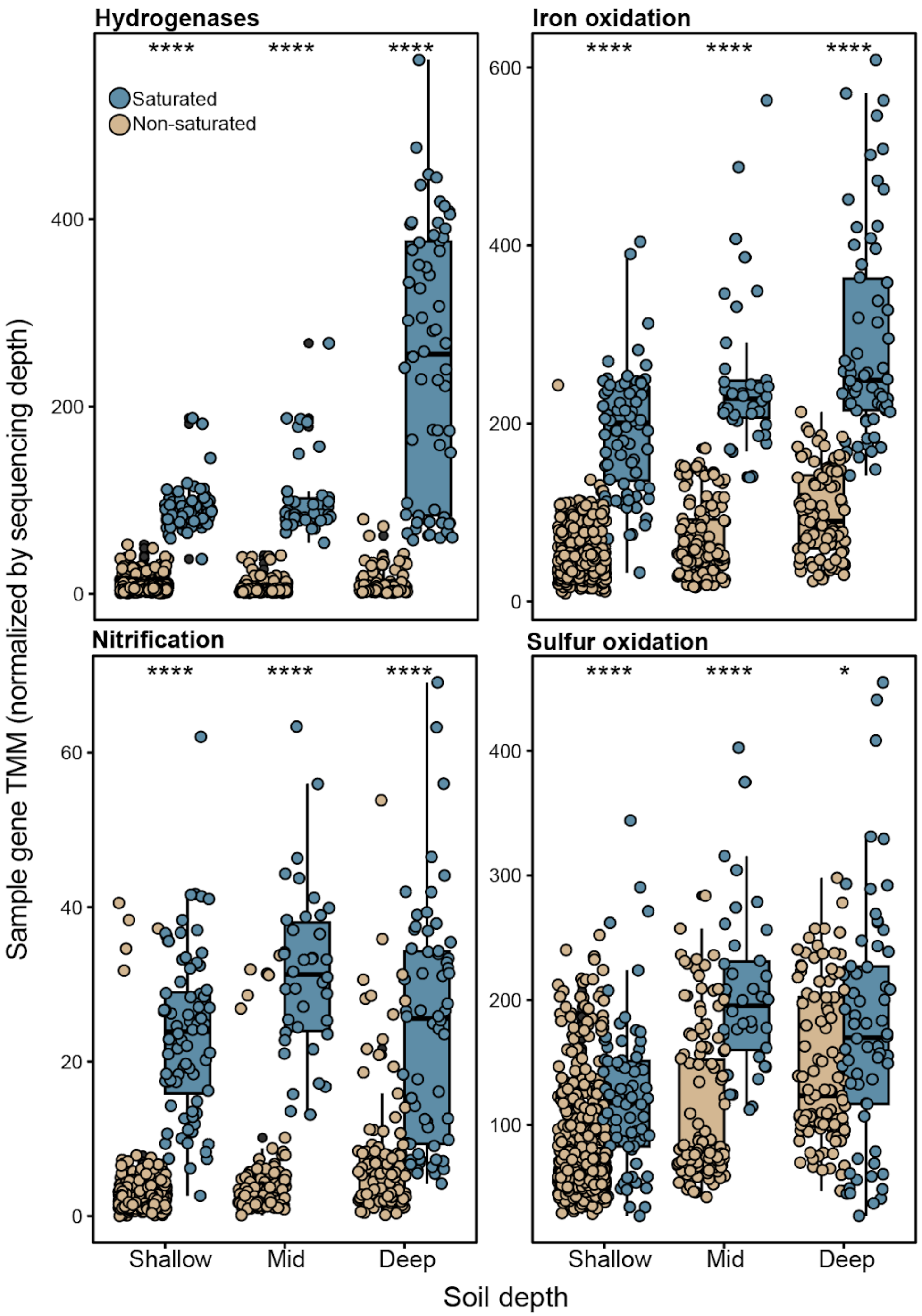


**Fig. S13.** Summed gene TMM of different metabolic or gene categories across bulk soil samples. Points represent individual soil samples. The lower and upper hinges of the boxplots represent the 25th and 75th percentiles, respectively, and the middle line is the median. The whiskers extend from the median by 1.5x the interquartile range. Points represent individual samples. Significant differences between saturated and non-saturated soils in each depth indicated with asterisks as indicated by Wilcoxon rank-sum test. *p<0.05, ****p<0.0001.

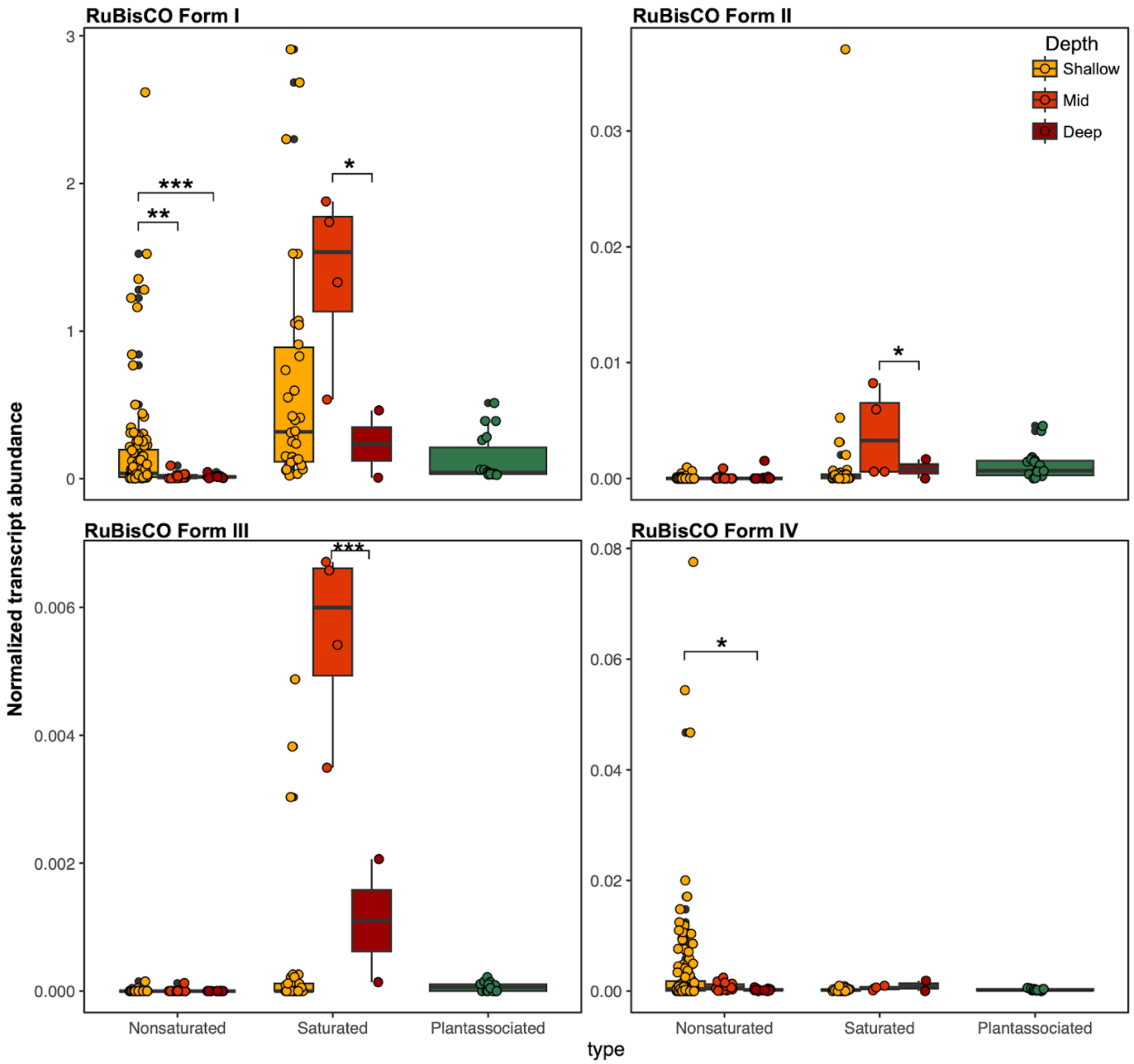


**Fig. S14.** Summed gene TMM of different RuBisCO forms across samples. Points represent individual soil samples. The lower and upper hinges of the boxplots represent the 25th and 75th percentiles, respectively, and the middle line is the median. The whiskers extend from the median by 1.5x the interquartile range. Points represent individual samples. Significant differences between soil depths indicated with asterisks as indicated by Wilcoxon rank-sum test. *p<0.05, **p<0.01, ***p<0.001, ****p<0.0001.

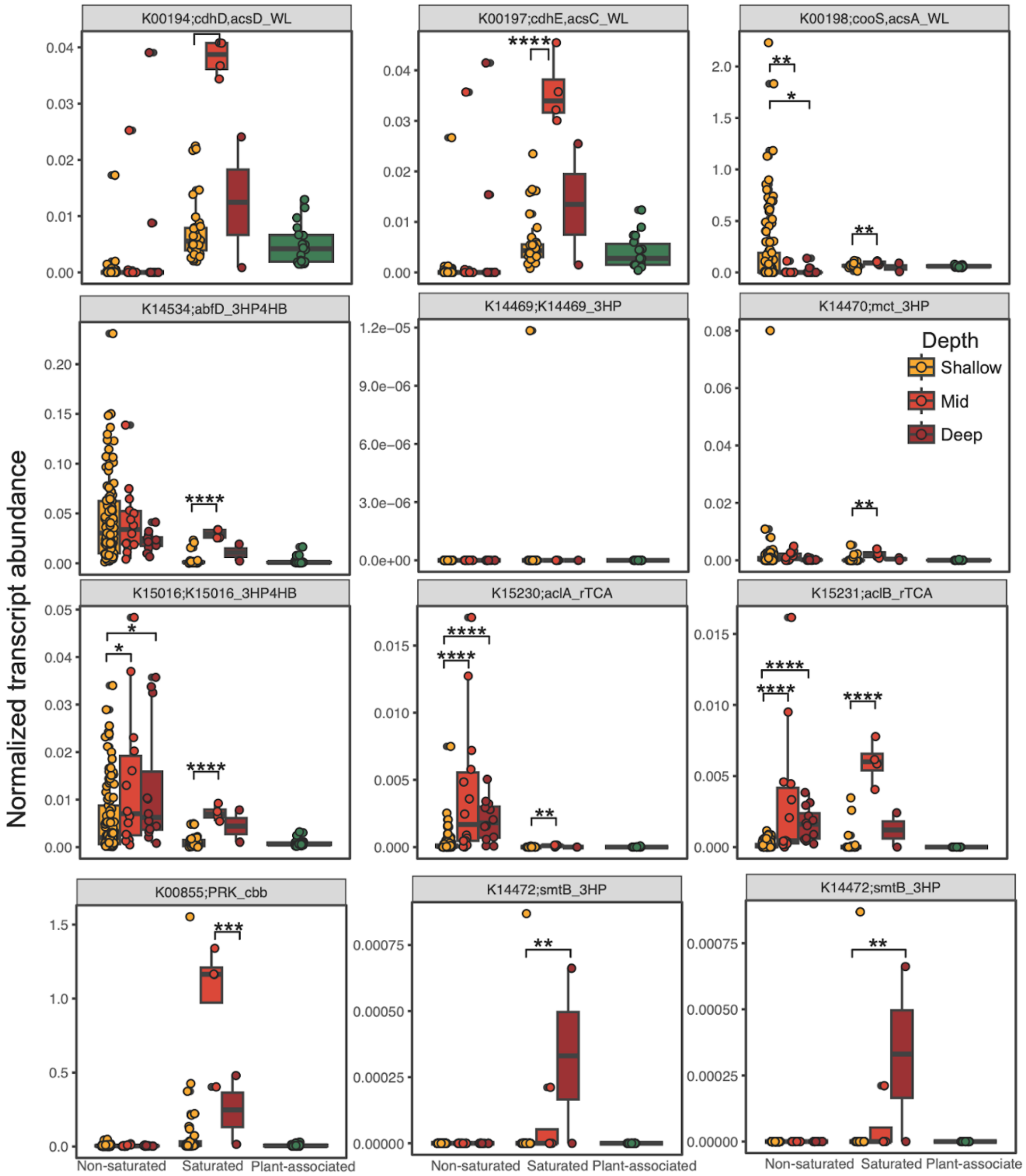


**Fig. S15.** Summed gene TMM of different autotrophy genes across samples. Points represent individual soil samples. The lower and upper hinges of the boxplots represent the 25th and 75th percentiles, respectively, and the middle line is the median. The whiskers extend from the median by 1.5x the interquartile range. Points represent individual samples. Significant differences between soil depths indicated with asterisks as indicated by Wilcoxon rank-sum test. *p<0.05, **p<0.01, ***p<0.001, ****p<0.0001.

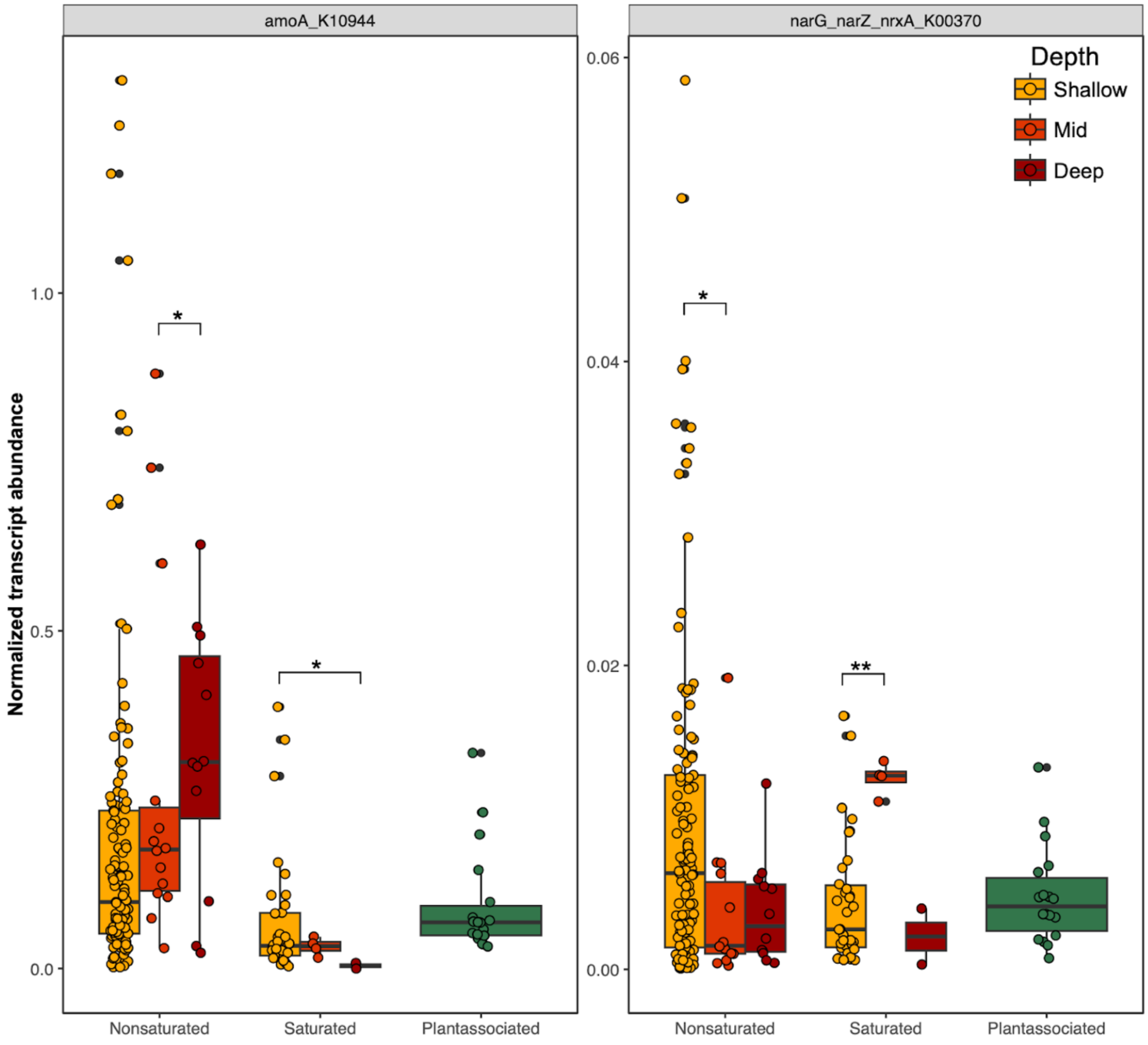


**Fig. S16.** Summed gene TMM of different nitrification genes across samples. Points represent individual soil samples. The lower and upper hinges of the boxplots represent the 25th and 75th percentiles, respectively, and the middle line is the median. The whiskers extend from the median by 1.5x the interquartile range. Points represent individual samples. Significant differences between soil depths indicated with asterisks as indicated by Wilcoxon rank-sum test. *p<0.05, **p<0.01, ***p<0.001, ****p<0.0001.

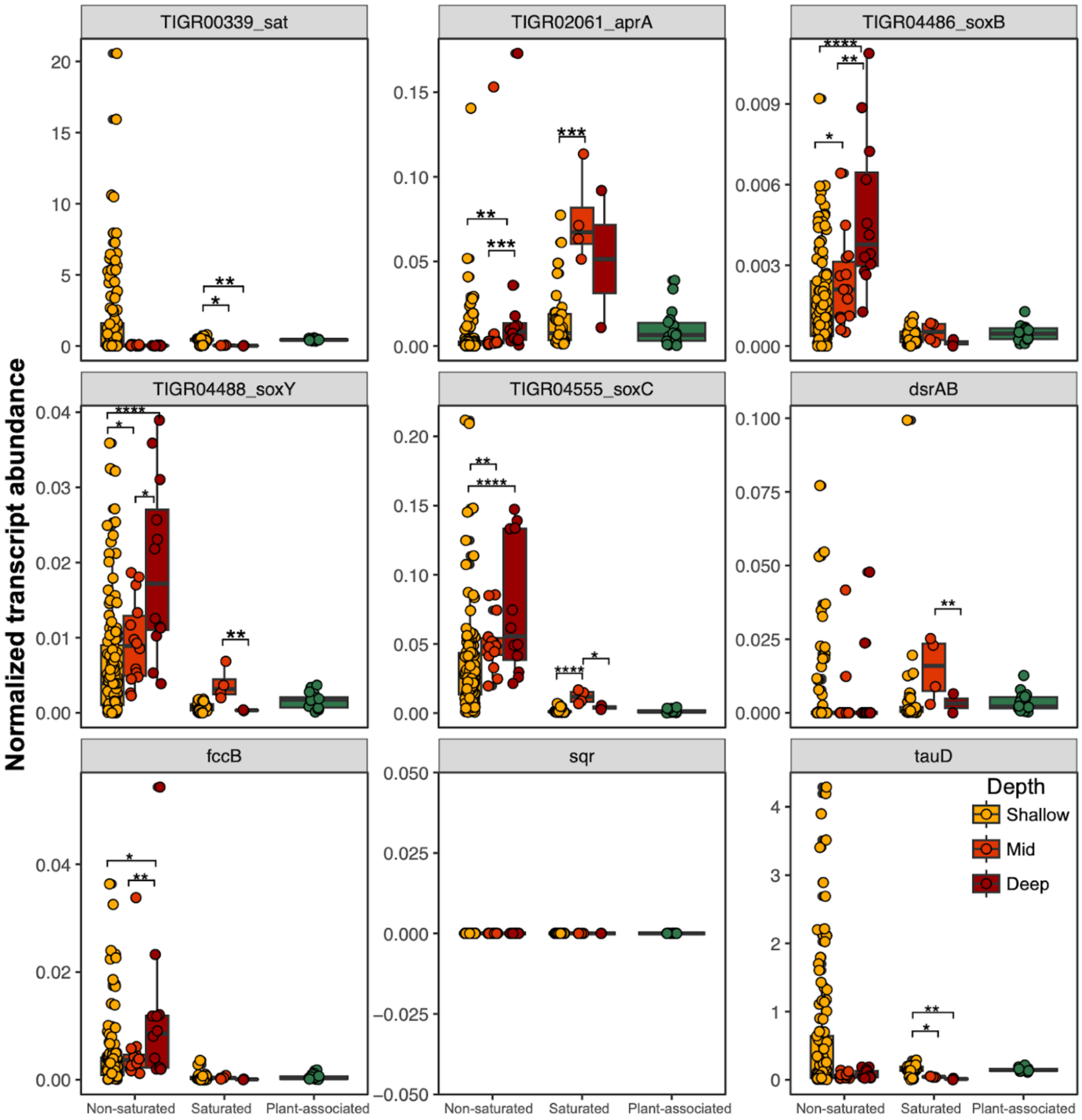


**Fig. S17.** Summed gene TMM of different sulfur oxidation genes across samples. Points represent individual soil samples. The lower and upper hinges of the boxplots represent the 25th and 75th percentiles, respectively, and the middle line is the median. The whiskers extend from the median by 1.5x the interquartile range. Points represent individual samples. Significant differences between soil depths indicated with asterisks as indicated by Wilcoxon rank-sum test. *p<0.05, **p<0.01, ***p<0.001, ****p<0.0001.

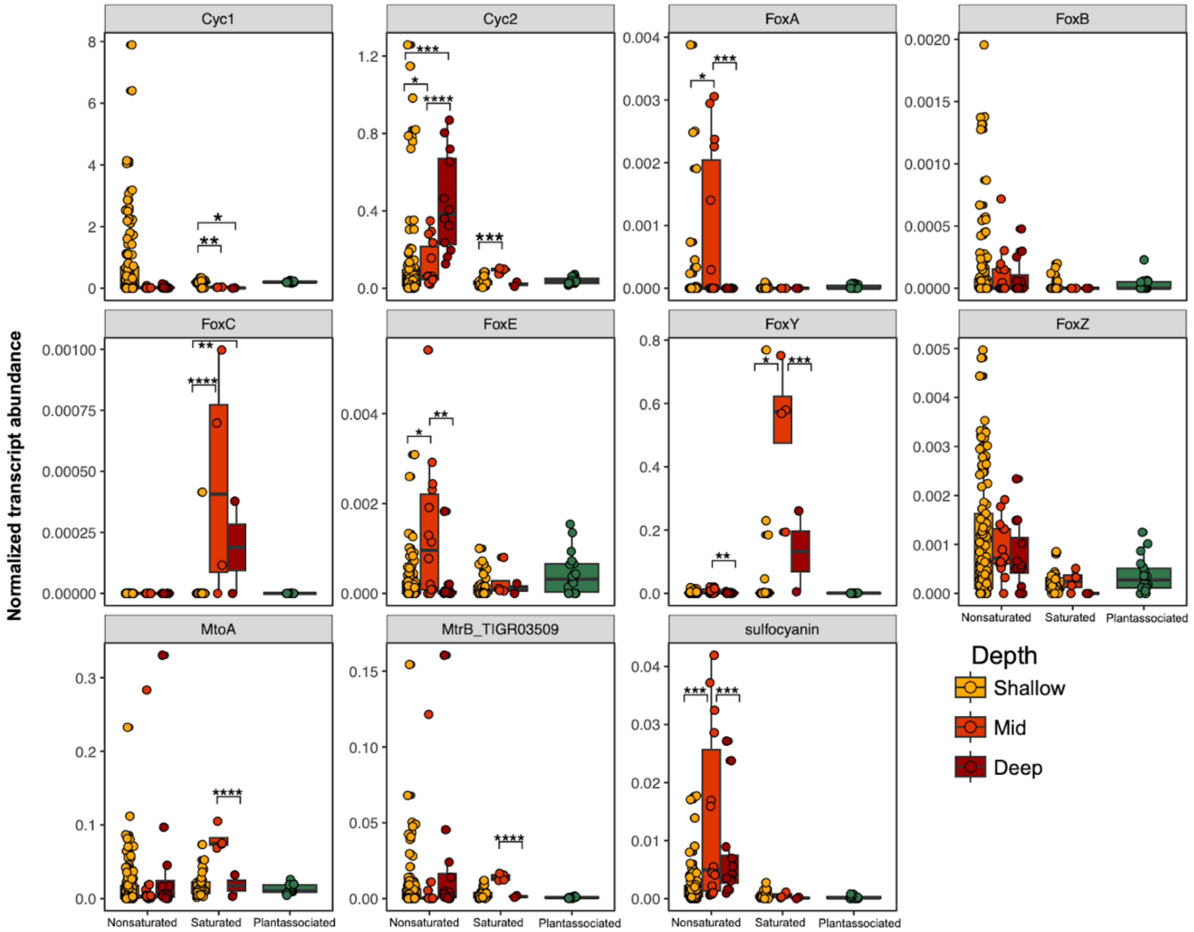


**Fig. S18.** Summed gene TMM of different iron oxidation genes across samples. Points represent individual soil samples. The lower and upper hinges of the boxplots represent the 25th and 75th percentiles, respectively, and the middle line is the median. The whiskers extend from the median by 1.5x the interquartile range. Points represent individual samples. Significant differences between soil depths indicated with asterisks as indicated by Wilcoxon rank-sum test. *p<0.05, **p<0.01, ***p<0.001, ****p<0.0001.

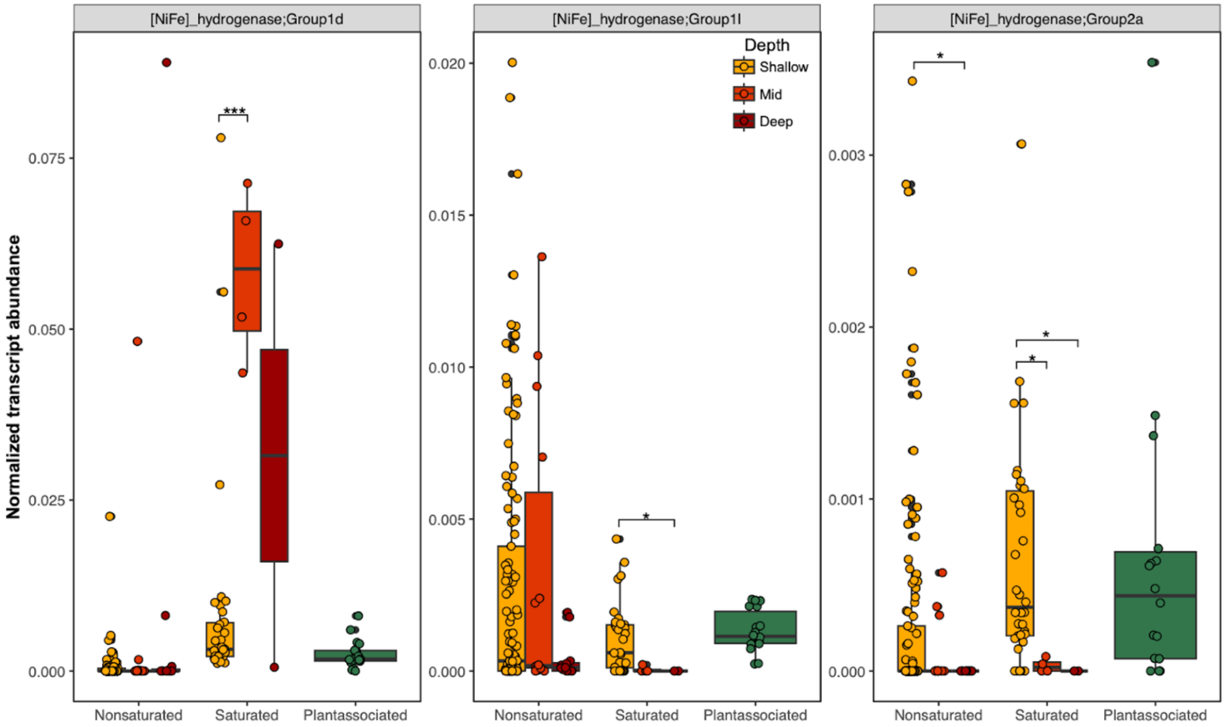


**Fig. S19.** Summed gene TMM of hydrogenases known to permit energy gain via aerobic respiration of $H_2$. Points represent individual soil samples. The lower and upper hinges of the boxplots represent the 25th and 75th percentiles, respectively, and the middle line is the median. The whiskers extend from the median by 1.5x the interquartile range. Points represent individual samples. Significant differences between soil depths indicated with asterisks as indicated by Wilcoxon rank-sum test. *p<0.05, **p<0.01, ***p<0.001, ****p<0.0001.

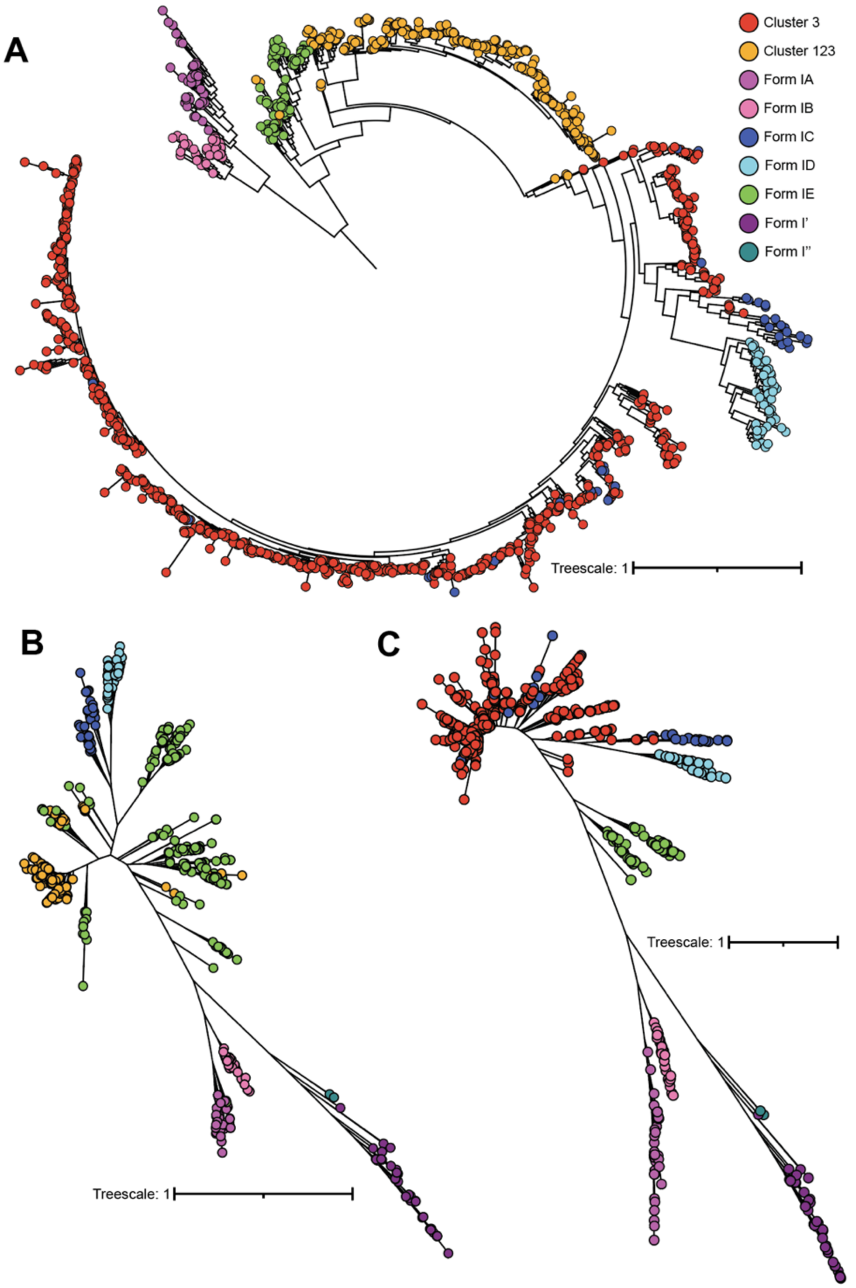


**Fig. S20. (A)** Concatenated RuBisCO protein phylogeny of all Cluster 3 and Cluster 123 sequences with Form I references from Prywes et al. (2023) and Ray et al. (2022) (1,2). Branches annotated with sequence origin (cluster number or reference annotation). **(B)** Concatenated RuBisCO protein phylogeny of all Cluster 123 sequences with Form I references. **(C)** Concatenated RuBisCO protein phylogeny of all Cluster 3 sequences with Form I references.

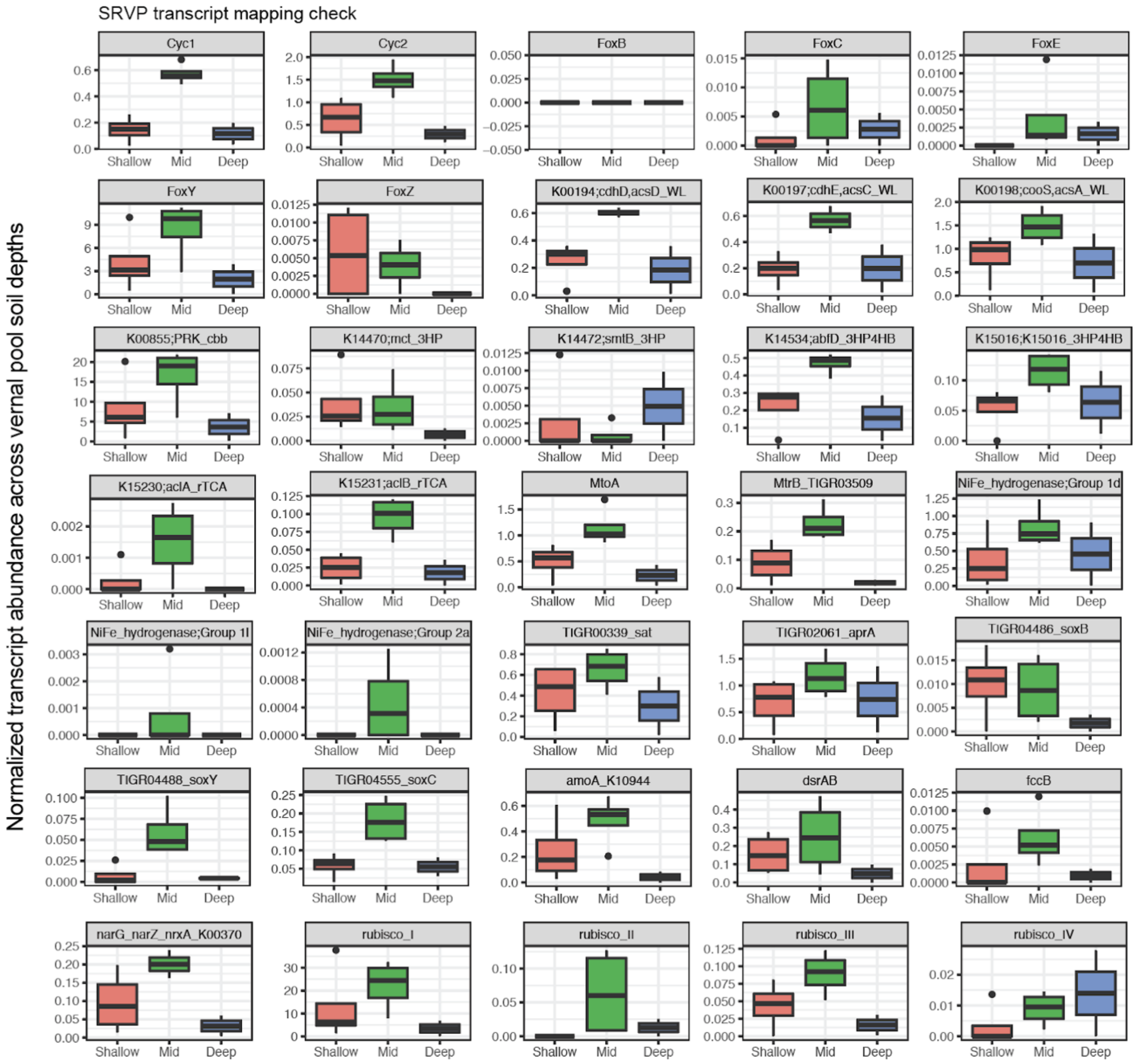


**Fig. S21.** Normalized transcript abundance of individual transcripts across the different soil depths of the vernal pools, generated by mapping metatranscriptomic reads to only genes recovered from vernal pool sediments. The lower and upper hinges of the boxplots represent the 25th and 75th percentiles, respectively, and the middle line is the median. The whiskers extend from the median by 1.5x the interquartile range.

**Supplementary tables**

| Ecosystem type/project | Shallow ≤25 cm | | Mid 25-50 cm | | Deep >50 cm | | Plant-associated | | Weathering granite | |
|---|---|---|---|---|---|---|---|---|---|---|
| Mixed-use agriculture | 6 | 0 | 6 | 0 | 0 | 0 | 0 | 0 | 0 | 0 |
| Colorado (subalpine floodplain) | 247 | 62 | 39 | 8 | 35 | 12 | 0 | 0 | 0 | 0 |
| Mediterranean grassland, Angelo | 62 | 8 | 66 | 6 | 33 | 0 | 27 | 0 | 0 | 0 |
| Mediterranean grassland, Hopland | 61 | 47 | 15 | 0 | 14 | 0 | 0 | 0 | 0 | 0 |
| Mediterranean grassland, Sagehorn | 4 | 0 | 4 | 0 | 16 | 0 | 0 | 0 | 0 | 0 |
| Rice paddy | 63 | 30 | 22 | 0 | 19 | 0 | 18 | 18 | 0 | 0 |
| Wetland | 20 | 4 | 17 | 4 | 41 | 2 | 0 | 0 | 0 | 0 |
| VIC, AUS (saprolite) | 0 | 0 | 0 | 0 | 0 | 0 | 0 | 0 | 18 | 0 |
| **Total condition metaG & metaT** | 463 | 151 | 169 | 18 | 158 | 14 | 45 | 18 | 18 | 0 |

**Table S1.** Number of metagenomes (white columns) and metatranscriptomes (grey columns) from each individual project, ecosystem type, and soil condition.

| Phyla | Form I | Form II | Form II/III | Form III | RLPs | Total |
|---|---|---|---|---|---|---|
| Proteobacteria | 1850 | 74 | 0 | 67 | 934 | 2925 |
| unknown | 794 | 20 | 4 | 197 | 587 | 1602 |
| Actinobacteria | 938 | 1 | 0 | 4 | 107 | 1050 |
| Chloroflexi | 179 | 4 | 0 | 18 | 46 | 247 |
| Planctomycetes | 21 | 1 | 0 | 6 | 177 | 205 |
| Firmicutes | 70 | 0 | 0 | 2 | 92 | 164 |
| Euryarchaeota | 39 | 5 | 0 | 70 | 28 | 142 |
| Bathyarchaeota | 40 | 4 | 0 | 56 | 36 | 136 |
| Bacteroidetes | 19 | 2 | 0 | 3 | 95 | 119 |
| Verrucomicrobia | 59 | 1 | 0 | 1 | 52 | 113 |
| Cyanobacteria | 82 | 0 | 0 | 0 | 9 | 91 |
| Armatimonadetes | 12 | 1 | 0 | 0 | 48 | 61 |
| NC10 | 39 | 0 | 0 | 0 | 7 | 46 |
| RIF-CHLX | 18 | 1 | 0 | 2 | 19 | 40 |
| Micrarchaeota | 10 | 0 | 0 | 3 | 6 | 19 |
| Nitrospirae | 7 | 0 | 0 | 1 | 10 | 18 |
| Acidobacteria | 5 | 0 | 0 | 1 | 10 | 16 |
| Elusimicrobia | 2 | 0 | 0 | 0 | 10 | 12 |
| Other (total <10) | 73 | 2 | 2 | 25 | 32 | 79 |

**Table S2.** Phyla assigned by ggKbase to RuBisCO protein sequences, across RuBisCO forms.

| Project or field site | Avg % transcripts mapping to genes from same site |
|---|---|
| Angelo | 93.5% |
| East River | 69.2% |
| Hopland | 61.8% |
| Rice paddies | 96.1% |
| Vernal pools | 98.3% |

**Table S3.** Average percentage of transcripts mapping to genes recovered from same project or field site.

**Supplementary Data File 1, Sheet A.** All sample information, metadata, and link to corresponding publications.

**Supplementary Data File 1, Sheet B.** List of all genes annotated, their assigned metabolic category, and annotation method (also outlined in methods).

**Supplementary Data File 1, Sheet C.** Final RuBisCO protein sequences, with origin sample, field site, soil depth, ecosystem type, assigned taxonomy.

**Supplementary Data File 1, Sheet D.** Final list of all annotated genes.

**Supplementary Data File 2.** Normalized coverage of all genes across samples (via metagenome read mapping to gene database).

**Supplementary Data File 3.** Normalized abundance of all transcripts across samples (via metatranscriptome read mapping to gene database).

### References

1. Tabita FR, Hanson TE, Li H, Satagopan S, Singh J, Chan S. Function, Structure, and Evolution of the RubisCO-Like Proteins and Their RubisCO Homologs. Microbiol Mol Biol Rev. 2007 Dec;71(4):576–99.

2. Prywes N, Phillips NR, Tuck OT, Valentin-Alvarado LE, Savage DF. Rubisco Function, Evolution, and Engineering. Annual Review of Biochemistry. 2023 June 20;92(Volume 92, 2023):385–410.

3. Ray AE, Zaugg J, Benaud N, Chelliah DS, Bay S, Wong HL, et al. Atmospheric chemosynthesis is phylogenetically and geographically widespread and contributes significantly to carbon fixation throughout cold deserts. The ISME Journal. 2022 Nov 1;16(11):2547–60.
